# XPA tumor variants lead to defects in NER that sensitize cells to cisplatin

**DOI:** 10.1101/2023.06.29.547124

**Authors:** Alexandra M. Blee, Kaitlyn S. Gallagher, Hyun-Suk Kim, Mihyun Kim, Christina R. Troll, Areetha D’Souza, Jiyoung Park, P. Drew Neufer, Orlando D. Schärer, Walter J. Chazin

## Abstract

Nucleotide excision repair (NER) neutralizes treatment with platinum (Pt)-based chemotherapy by removing Pt lesions from DNA. Previous study has identified that missense mutation or loss of either of the NER genes Excision Repair Cross Complementation Group 1 and 2 (*ERCC1* and *ERCC2*) leads to improved patient outcomes after treatment with Pt-based chemotherapies. Although most NER gene alterations found in patient tumors are missense mutations, the impact of such mutations in the remaining nearly 20 NER genes is unknown. Towards this goal, we previously developed a machine learning strategy to predict genetic variants in an essential NER scaffold protein, Xeroderma Pigmentosum Complementation Group A (XPA), that disrupt repair activity on a UV-damaged substrate. In this study, we report in-depth analyses of a subset of the predicted NER-deficient XPA variants, including *in vitro* analyses of purified recombinant protein and cell-based assays to test Pt agent sensitivity in cells and determine mechanisms of NER dysfunction. The most NER deficient variant Y148D had reduced protein stability, weaker DNA binding, disrupted recruitment to damage, and degradation resulting from tumor missense mutation. Our findings demonstrate that tumor mutations in XPA impact cell survival after cisplatin treatment and provide valuable mechanistic insights to further improve variant effect prediction efforts. More broadly, these findings suggest XPA tumor variants should be considered when predicting patient response to Pt-based chemotherapy.

**Significance:** A destabilized, readily degraded tumor variant identified in the NER scaffold protein XPA sensitizes cells to cisplatin, suggesting that XPA variants can be used to predict response to chemotherapy.

## Introduction

Nucleotide excision repair (NER) protects cells from DNA damage by removing bulky DNA adducts such as those introduced by car exhaust, cigarette smoke, ultraviolet (UV) light, and platinum (Pt)-based chemotherapeutics. (1) Germline mutations in genes that reduce NER activity cause the inherited disorder *xeroderma pigmentosum* (XP), hallmarked by extreme sensitivity to solar UV radiation and an over 2000-fold increased incidence of skin cancer. (2–4) Reduced NER activity is also correlated with improved tumor response to Pt chemotherapy.

Nonrecurrent somatic missense mutations in Excision Repair Cross Complementation Group 2 (*ERCC2*) that lead to defective NER or loss of *ERCC1* has been shown to sensitize tumor cells to cisplatin, leading to improved patient outcomes. (5–8) However, the impact of genetic alterations in the remaining NER genes toward Pt therapy remains unknown. Further investigation is motivated by the study of The Cancer Genome Atlas (TCGA) Pan-Cancer Atlas, which revealed that most genetic alterations in NER genes are nonrecurrent nonsynonymous single nucleotide variants of unknown significance. (9) The ability to accurately identify deleterious mutations in NER genes from this pool and to determine the mechanisms of reduced repair efficiency that sensitize cells to Pt drugs would better inform clinical decision-making and guide selection of the most successful therapy for individual patients.

Sequence-based genetic variant interpretation represents one strategy to identify mutations that will impact tumor response to therapy, but is severely limited by insufficient benchmarking, lack of data from diverse populations, and absence of functional validation. (10) As a robust alternative, we have previously developed a machine learning approach that incorporates functional validation to more accurately predict variants that cause NER defects in cells. (11) This prior study focused on tumor variants in the essential NER scaffold protein, Xeroderma Pigmentosum Complementation Group A (XPA), and identified a set of variants predicted to sensitize cells to cisplatin. However, the functional analyses were limited to a single DNA repair assay. Here, we report a thorough analysis of a subset of these mutants characterizing their physical properties, NER capacity, and sensitivity to both UV and cisplatin. This investigation is designed to (1) provide evidence to test the hypothesis that NER tumor variants contribute to drug sensitivity in cells, (2) provide proof-of-concept that our machine learning strategy can predict tumor variants that cause functional repair defects and sensitize cells to cisplatin, and (3) elucidate mechanisms of NER dysfunction.

Five XPA variants from our previous study (F1112C, M113I, D114Y, R130I, Y148D) were of particular interest because they have relatively high confidence predictions of NER-deficiency (11) and are all located in functionally-relevant regions of the globular DNA binding domain (DBD) of XPA. Interestingly, of these five, only M113I, D114Y, and Y148D exhibited NER defects in the repair of a UV-damaged reporter using the fluorescence multiplex host cell reactivation (FM-HCR) functional validation assay. (11) These discrepancies between the machine learning predictions and repair activity in cells motivated further study. In addition, characterization of these variants provided an opportunity to test repair activity on cisplatin lesions and identify mechanisms of dysfunction associated with each by assessing variant protein structure and stability, DNA binding affinity, interaction with RPA, and recruitment to sites of damage. *In vitro* analyses of purified recombinant variant protein and cell-based assays revealed that our established machine learning strategy can identify XPA variants that disrupt NER and sensitize cells to cisplatin. It also revealed XPA variant protein destabilization, disrupted DNA binding and recruitment to damage, and degradation in cells as key mechanisms of NER dysfunction. The results are discussed in the context of the ongoing efforts to reliably identify tumor variants that will impact response to Pt-based chemotherapeutics.

## Materials and Methods

### Cell Lines and Cell Culture

SV40-transformed human dermal fibroblast XP2OS (XPA-null) cells (12) (RRID:CVCL_F510) were reported in a recent publication. (13) SV40-transformed HEK293FT cells (RRID:CVCL_6911) were gifted from Drs. Margaret Axelrod and Justin Balko (Vanderbilt University Medical Center). Cells were cultured in Dulbecco’s Modified Eagle’s Medium (DMEM) with high glucose, pyruvate, and GlutaMAX (Thermo Fisher Scientific #10569044) supplemented with 10% qualified One Shot United States Origin Fetal Bovine Serum (FBS) (Thermo Fisher Scientific #A3160502) and 1% Penicillin-Streptomycin (Thermo Fisher Scientific #15140122) at 37 °C with 5% CO2, unless indicated otherwise. Cells were used within three months of thawing and passage number did not exceed 35 for any experiments. Cells were routinely screened and confirmed negative for mycoplasma contamination at least every three months (SouthernBiotech #13100-01).

## Plasmids

For recombinant protein expression of both full-length (residues 1-239) and DNA binding domain (residues 98-239) human XPA, cDNA (NM_000380) was cloned into the pBG100 vector (RRID:Addgene_33365) with an N-terminal His_6_ tag. Q5 site-directed mutagenesis (New England BioLabs #E05545) was used to generate F112C, M113I, D114Y, R130I, or Y148D XPA variants. Mutagenesis primers are provided in the supplementary information as **Supplementary Table 1**.

For recombinant protein expression of full-length RPA protein, codon optimized RPA70 was cloned into the pBAD3C vector and RPA14 and 32 were cloned into the pRSFDuet3C vector.

For lentiviral transduction, the pMD2.G (RRID:Addgene_12259), psPAX2 (RRID:Addgene_12260), and pWPXL (RRID:Addgene_12257) were used as described. (13) Full-length wild-type (WT), F112C, M113I, D114Y, R130I, or Y148D XPA cDNA (NM_000380) was inserted into pWPXL from the previously generated pBG100 (RRID:Addgene_33365) vectors using restriction enzyme digest. Additional plasmids for recombinant expression of the remaining NER proteins besides XPA and RPA (XPC-RAD23B, (14) XPG, (15) XPF-ERCC1, (16) and TFIIH (17)) have been reported previously.

## Antibodies

The following antibodies were used for Western blotting: rabbit polyclonal anti-XPA (Abcam ab85914, RRID:AB_1925572) diluted 1:1,000, rabbit monoclonal anti-β-tubulin (Cell Signaling Technology #2128S) diluted 1:1,000, rabbit monoclonal anti-ubiquitin Lys48-specific (Millipore Sigma 05-1307, RRID:AB_1587578) diluted 1:1,000, goat anti-rabbit IgG (H+L) HRP conjugate (Millipore Sigma #AP307P, RRID:AB_92641) diluted 1:5,000; and for immunofluorescence: rabbit polyclonal anti-XPA (Abcam ab85914, RRID:AB_1925572) diluted 1:5,000, mouse monoclonal anti-(6–4) Photoproducts (6-4PP, Cosmo Bio LTD #CAC-NM-DND-002, RRID:AB_1962842) diluted 1:400, goat anti-rabbit IgG (H+L) Alexa Fluor 488 (Thermo Fisher Scientific #A11008, RRID:AB_143165) diluted 1:1,000, and goat anti-mouse IgG (H+L) Alexa Fluor 594 (Thermo Fisher Scientific #A11032, RRID:AB_2534091) diluted 1:1,000.

## Lentiviral Cell Line Transduction

Lentiviral particles were generated in HEK293FT cells (RRID:CVCL_6911) in 6-well dishes at 70-80% confluency at time of transfection, as described previously. (13) Cells were transfected in lentivirus packaging medium containing Opti-MEM supplemented with 5% FBS (Thermo Fisher Scientific #A3160502) and 1 mM sodium pyruvate (Thermo Fisher Scientific #11360070). Transfectant solution included 2.25 μg pMD2.G (RRID:Addgene_12259), 2.25 μg psPAX2 (RRID:Addgene_12260), and 0.75 μg pWPXL (RRID:Addgene_12257) containing WT, F112C, M113I, D114Y, R130I, or Y148D XPA, or a mock transfection of XPA-null cells, using 7 μL Lipofectamine 3000 and 6 μL P3000 (Thermo Fisher Scientific #L3000008) in 500 μL total Opti-MEM reduced serum media with GlutaMAX (Thermo Fisher Scientific #51985034) and following manufacturer’s instructions for improved lentiviral production. Target XPA-null XP2OS cells (RRID:CVCL_F510) were seeded in 6-well dishes and allowed to reach 50% confluency by time of transduction. Viral particle-containing supernatant was collected 24-and 52-hours post-transfection from HEK293FT cell (RRID:CVCL_6911) plates, 0.2 μm-filtered, and used to transduce XPA-null XP2OS cells (RRID:CVCL_F510). Transduced XP2OS cells (RRID:CVCL_F510) were subsequently cultured as described in the Cell Lines and Cell Culture section.

## Western Blotting

Cell pellets were harvested on ice and washed with Dulbecco’s Phosphate Buffered Saline (DPBS) without calcium chloride or magnesium chloride (Thermo Fisher Scientific #14190-144) before lysing in RIPA lysis buffer (150 mM NaCl, 5 mM EDTA, 50 mM Tris, 1% NP-40, 0.5% sodium deoxycholate, 0.1% SDS, pH 8.0) supplemented with 1X protease inhibitor cocktail (Millipore Sigma #P8340). Lysates were centrifuged to remove cellular debris and protein concentration was measured by DC Protein Concentration Assay (Bio-Rad #5000112) following manufacturer’s instructions. Equal amounts of protein (∼50 μg) from cell lysates were denatured in 1X NuPAGE LDS sample buffer (Thermo Fisher Scientific #NP0007) containing 0.1 mM DTT and RIPA buffer, and heated at 95 °C for 5 minutes. Samples were subjected to SDS-polyacrylamide gel electrophoresis (PAGE) in 4-12% NuPAGE Bis-Tris gels (Thermo Fisher Scientific #NP0326BOX) at 100 V for 90 minutes and wet-transferred to nitrocellulose membranes (Thermo Fisher Scientific #88018) at 30 V for 90 minutes following manufacturer’s instructions. Membranes were briefly stained with Ponceau S (Millipore Sigma #7170) to check for successful transfer. Membranes were blocked in 5% w/v milk in 1X Tris-buffered saline-Tween 20 (TBST, 20 mM Tris, 150 mM NaCl, 0.1% w/v Tween 20, pH 7.6) for 1 hour at room temperature and immunoblotted with indicated primary antibodies overnight at 4 °C diluted 1:1000 in 5% w/v bovine serum albumin (BSA) in 1X TBST. Membranes were washed 3 times 5 minutes per wash in 1X TBST at room temperature and incubated with horseradish peroxidase (HRP)-conjugated secondary antibodies diluted 1:5000 in 5% milk w/v in 1X TBST for one hour at room temperature. Blots were visualized after incubation for 2 minutes with Pierce ECL Western Blotting Substrate (Thermo Fisher Scientific #32209) following manufacturer’s instructions and imaged using a Amersham Imager 600 (GElifesciences/Cytiva). All blot images for analysis were collected prior to detector saturation.

## RT-qPCR

Cell pellets were harvested on ice, washed with 1X DPBS (Thermo Fisher Scientific #14190-144), and lysed in TRIzol (Thermo Fisher Scientific #15596026). RNA was extracted from the TRIzol samples following manufacturer’s instructions, using chloroform to separate RNA from DNA and proteins, isopropanol to precipitate the RNA, 75% ethanol to wash the RNA, and nuclease-free water (Thermo Fisher Scientific #10977015) to resuspend the purified RNA pellet. Reverse transcription was performed using the SuperScript IV VILO Master Mix with ezDNase Enzyme reverse transcription kit (Thermo Fisher Scientific #11766050) following manufacturer’s instructions including DNase treatment prior to reverse transcription. cDNA was amplified in a CFX96 Touch Real-Time PCR Detection System (Bio-Rad) using Taqman Gene Expression Assay (FAM) probes for *GAPDH* (Thermo Fisher Scientific #4331182, ID: Hs02786624_g1) and *XPA* (Thermo Fisher Scientific #4331182, ID: Hs00902270_m1) following manufacturer’s instructions. All reactions for each independent replicate were performed using 100 ng cDNA in triplicate. Relative fold difference in transcript abundance was determined using the ΔΔC_t_ method. (18)

## Clonogenic Cell Survival Assay with Ultraviolet (UV)-C Irradiation

Cells were plated at single cell density in 6 cm dishes with 275 cells per well and clonogenic cell survival was assayed similarly to as described previously. (5,19–21) Cells were allowed to adhere overnight following standard culture conditions, washed with 1X DPBS (Thermo Fisher Scientific #14190-144), and exposed to indicated dose of UV-C (254 nm) radiation before the culture medium was replenished and cells were returned to the temperature and CO_2_-controlled incubator. 14 days after exposure, cells were fixed with 4% paraformaldehyde (Fisher Scientific #50-980-487) diluted in 1X PBS (137 mM NaCl, 2.7 mM KCl, 10 mM Na_2_HPO_4_, 1.8 mM KH_2_PO_4_, pH 7.4) for 15 minutes at room temperature. The fixative was removed and cells were washed with 1X PBS and then stained with 0.5% crystal violet (Fisher Scientific #C581-25) diluted in water for 60 minutes. Excess stain was removed and dishes were washed with water. Dishes were imaged using a E-Gel Imager (Thermo Fisher Scientific). Colonies (defined as a minimum of 50 cells) were counted using FIJI. Percent survival was calculated as the percentage of colonies that grew on the treated dishes relative to the untreated dishes at each indicated dose.

## Sulforhodamine B Colorimetric Assay

To test sensitivity to cisplatin, for each independent replicate, cells were plated in triplicate at a density of 4 x 10^4^ cells per well for each planned genotoxin dose, using 96-well plates. Cells were allowed to adhere overnight before treatment with indicated genotoxin following standard culture conditions, incubation for 120 hours, fixation, and staining as reported previously. (22–24) At time of treatment, standard culture media was removed and replaced with serum-and antibiotic-free media containing the indicated concentration of genotoxin (zero hour timepoint) and cells were incubated under standard culture conditions for 2 hours. After incubation, genotoxin-containing media was removed, wells were washed with 1X DPBS (Thermo Fisher Scientific #14190-144), and standard culture media was replaced. Cells were cultured for an additional 118 hours under standard culture conditions. At the 120-hour endpoint, cells were fixed by removing standard culture media and adding 10% w/v trichloroacetic acid (TCA, Millipore Sigma #91228) and incubation at 4 °C for 60 minutes. Excess 10% TCA was removed, plates were washed by dipping into a basin of gently running water four times, and dried at room temperature. Cells were stained using 0.4% w/v sulforhodamine B sodium salt (Millipore Sigma #S1402) in 1% acetic acid for 30 minutes at room temperature. Excess stain was removed, each well was washed with 200 μL of 1% acetic acid four times, and plates were dried at room temperature. Protein-bound sulforhodamine B dye was solubilized by adding 100 μL of 10 mM Tris base (Fisher Scientific #BP152-500) to each well and incubation with shaking for 10 minutes. Absorbance of each well at 490 nm was measured using a BioTek Synergy H1 Hybrid Reader microplate reader. For each independent replicate, a fresh stock of *cis*-Diamineplatinum(II) dichloride (cisplatin, Millipore Sigma, #479306) stock was made by diluting to 2 mM in 0.09% NaCl and incubated for 12 hours at room temperature with gentle shaking prior to use, following manufacturer’s instructions. To calculate the percent of untreated cell growth, the mean absorbance values were calculated for each triplicate condition and subtracted by the mean absorbance from media-only control wells. The media-subtracted mean absorbances of each genotoxin concentration were divided by those for the corresponding untreated (zero hour) wells and multiplied by 100.

To test proliferation in untreated cells, for each independent replicate, cells were plated in triplicate at a density of 4 x 10^4^ cells per well for each cell line and timepoint, using 96-well plates. Cells were incubated following standard culture conditions for indicated timepoints, fixed, and stained as described following the same method as for cisplatin sensitivity.

## Bortezomib Treatment

For each independent replicate, cells were plated at a density of 1 x 10^6^ cells per well of a 6-well plate and allowed to adhere overnight following standard culture conditions. To inhibit the Ubiquitin Proteasome System (UPS), cells were incubated with 100 nM bortezomib (Fisher Scientific #50-187-1974) for either 0, 2, 4, and 8 hours in standard culture media before harvest and processing as described in the Western Blotting section.

## XPA Expression and Purification

Full-length WT or variant XPA protein with a N-terminal His_6_ tag in the pBG100 vector was expressed in *Escherichia coli* Rosetta2 pLysS competent cells grown in Terrific Broth medium with 50 μg/mL kanamycin and 10 μM ZnCl_2_, and cell growth and protein purification were performed similarly to our previous study. (13) Cells were grown to OD_600_ = 0.6 at 37 °C and then to OD_600_ = 1.2 at 18 °C, at which point protein expression was induced with 0.5 mM isopropyl β-D-1-thiogalactopyranoside (IPTG) and cells were grown at 18 °C for 16 hours. Cells were harvested by centrifugation at 6,500 rpm for 20 minutes at 4 °C. All subsequent purification steps were performed on ice or at 4 °C unless indicated. Cell pellets were resuspended in 10 mL of Lysis Buffer per 1 g of pellet (100 mM Tris pH 8.0, 500 mM NaCl, 20 mM imidazole, 10% glycerol, 5 mM β-mercaptoethanol, 200 μg/mL lysozyme, 10 μL of Roche DNase I recombinant (Millipore Sigma #04536282001), 5 mM magnesium acetate, 1 Roche cOmplete EDTA-free Protease Inhibitor Cocktail tablet (Millipore Sigma #04693132001), 0.5 mM phenylmethylsulfonyl fluoride (PMSF), 1 mM benzamidine) and Dounce homogenized. Cells were further lysed by sonication at 60% amplitude (5 seconds on/10 seconds off) for 10 minutes. Lysate was clarified by centrifugation at 20,000 rpm for 40 minutes at 4 °C and filtration through a 0.45 μm polyvinylidene difluoride (PVDF) syringe filter. Supernatant was incubated with the equivalent of a 5 mL bed of Ni Sepharose High Performance resin (Millipore Sigma #GE17-5268-01) equilibrated with Ni-NTA Buffer A (100 mM Tris pH 8.0, 500 mM NaCl, 20 mM imidazole, 10% glycerol, 5 mM β-mercaptoethanol) for 60 minutes before passing over a gravity column. Resin was washed with 10 column volumes (CV) of Ni-NTA Buffer A and protein was eluted Ni-NTA Buffer B (100 mM Tris pH 8.0, 500 mM NaCl, 400 mM imidazole, 10% glycerol, 5 mM β-mercaptoethanol). Eluent was incubated with H3C protease for 16 hours to cleave the His_6_ tag and dialyzed in Pre-Heparin Buffer (20 mM Tris pH 8.0, 300 mM NaCl, 1 mM DTT, 10% glycerol). Dialyzed, cleaved protein was diluted to a final salt concentration of 150 mM NaCl using Dilution Buffer (20 mM Tris pH 8.0, 1 mM DTT, 10% glycerol), applied to a 5 mL HiTrap Heparin HP column (Cytiva #17040701) equilibrated with Heparin Buffer A (20 mM Tris pH 8.0, 50 mM NaCl, 10% glycerol, 1 mM DTT), washed with 5 CV Heparin Buffer A, washed with a linear gradient of 0-30% Heparin Buffer B (20 mM Tris pH 8.0, 1 M NaCl, 10% glycerol, 1 mM DTT) over 8 CV, and eluted using a linear gradient of 30-70% Heparin Buffer B over 12 CV followed by 5 CV of 100% Heparin Buffer B. XPA protein eluted at approximately 45% Heparin Buffer B. Eluted protein was further purified on a 120 mL HiLoad 16/600 Superdex 75 pg column (Cytiva #28-9893-33) equilibrated with S75 Buffer (50 mM Tris pH 8.0, 150 mM NaCl, 10% glycerol, 1 mM DTT). XPA protein eluted at approximately 65 mL. Protein identity was confirmed by electrospray ionization mass spectrometry and used for experimentation within 24 hours after final elution, or aliquoted and flash frozen in liquid nitrogen.

WT or variant XPA DNA binding domain (DBD) protein (residues 98-239) with a N-terminal His_6_ tag in the pBG100 vector was expressed in *Escherichia coli* BL21 DE3 competent cells grown in Terrific Broth medium with 50 μg/mL kanamycin and 10 μM ZnCl_2_. Cell growth and harvest was performed following the same method as for full-length XPA. All subsequent purification steps were performed on ice or at 4 °C unless indicated. Cell pellets were resuspended in 10 mL of Ni-NTA Buffer A per 1 g of pellet (20 mM Tris pH 7.5, 500 mM NaCl, 15 mM imidazole, 10% glycerol) plus 1 Roche cOmplete EDTA-free Protease Inhibitor Cocktail tablet (Millipore Sigma #04693132001), 0.5 mM PMSF, and 1 mM benzamidine. Cells were lysed by sonication at 50% amplitude (5 seconds on/10 seconds off) for 10 minutes. Lysate was clarified by centrifugation at 20,000 rpm for 20 minutes at 4 °C and filtration through a 0.45 μm PVDF syringe filter.

Supernatant was incubated with the equivalent of a 10 mL bed of Ni Sepharose High Performance resin (Millipore Sigma #GE17-5268-01) equilibrated with Ni-NTA Buffer A (100 mM Tris pH 8.0, 500 mM NaCl, 20 mM imidazole, 10% glycerol, 5 mM β-mercaptoethanol) for 60 minutes before passing over a gravity column. Resin was washed with 10 column volumes (CV) of Ni-NTA Buffer A and protein was eluted Ni-NTA Buffer B (20 mM Tris pH 7.5, 500 mM NaCl, 400 mM imidazole, 10% glycerol). Eluent was diluted to a final salt concentration of 150 mM NaCl using Dilution Buffer (20 mM Tris pH 7.5, 1 mM DTT, 10% glycerol) and incubated with H3C protease for 3 hours to cleave the His_6_ tag. Diluted, cleaved protein was applied to a 5 mL HiTrap Heparin HP column (Cytiva #17040701) equilibrated with Heparin Buffer A (20 mM Tris pH 7.5, 150 mM NaCl, 10% glycerol, 1 mM DTT), washed with 8 CV Heparin Buffer A, and eluted using a linear gradient of 0-100% Heparin Buffer B (20 mM Tris pH 8.0, 1 M NaCl, 10% glycerol, 1 mM DTT) over 12 CV followed by 3 CV of 100% Heparin Buffer B. XPA protein eluted at approximately 50% Heparin Buffer B. Eluted protein was further purified on a 24 mL Superdex Increase 75 10/300 column (Cytiva #29-1487-21) equilibrated with S75 Buffer (20 mM Tris pH 7.5, 150 mM NaCl, 10% glycerol, 1 mM DTT). XPA protein eluted at approximately 11 mL. Protein identity was confirmed by electrospray ionization mass spectrometry and used for experimentation within 24 hours after final elution, or aliquoted and flash frozen in liquid nitrogen.

For both the full-length and DBD constructs of the Y148D XPA variant, refolding was necessary to obtain yields of soluble protein. The expression and purification methods were modified as follows. Cells were grown in Luria-Bertani (LB) broth medium with 50 μg/mL kanamycin and 10 μM ZnCl_2_. Cells were grown to OD_600_ = 0.6 at 37 °C and then to OD_600_ = 1.2 at 18 °C, at which point protein expression was induced with 0.25 mM IPTG and cells were grown at 18 °C for 16 hours. Cells were harvested by centrifugation at 6,500 rpm for 20 minutes at 4 °C. All subsequent purification steps were performed on ice or at 4 °C unless indicated. Cell pellets were resuspended and lysed following the same method as described for WT and other variant XPA proteins. Lysate was clarified by centrifugation at 20,000 rpm for 20 minutes at 4 °C. Insoluble Y148D XPA protein present in the pellet after centrifugation was subjected to a second round of resuspension, lysis, and centrifugation as described. Taking the pelleted sample, resuspend in 10 mL of Ni-NTA Buffer A with 4 M guanidinium hydrochloride (GuHCl) per original 1 g of original pellet. Cells were lysed by sonication at 50% amplitude (5 seconds on/10 seconds off) for 10 minutes. Lysate was clarified by centrifugation at 20,000 rpm for 20 minutes at 4 °C. The denatured protein in the supernatant was collected and four serial dialysis steps into Ni-NTA Buffer A were performed to remove the GuHCl from the sample. Dialyzed sample was filtered through a 0.45 μm PVDF syringe filter. Purification continued beginning with Ni Sepharose High Performance resin as described for WT and other XPA variant proteins.

## RPA Expression and Purification

Full-length RPA protein (RPA70, RPA32, and RPA14 constructs) was expressed in Rosetta2 pLysS competent cells grown in Terrific Broth medium with 50 μg/mL kanamycin and 100 μg/mL ampicillin. Cells were grown to OD_600_ = 0.8 at 37 °C and then to OD_600_ = 1.1 at 18 °C, at which point protein expression was induced with 1 mM IPTG and 2 g/L of L-Arabinose and cells were grown at 18 °C for 16 hours. Cells were harvested by centrifugation at 6,500 rpm for 20 minutes at 4 °C. All subsequent purification steps were performed on ice or at 4 °C unless indicated. Cell pellets were resuspended in 5 mL of Lysis Buffer per 1 g of pellet (20 mM HEPES pH 7.5, 500 mM NaCl, 5 mM β-mercaptoethanol, 10 μM ZnCl_2_, 10 mM imidazole, 2 Roche cOmplete EDTA-free Protease Inhibitor Cocktail tablets (Millipore Sigma #04693132001). Cells were further lysed by sonication at 60% amplitude (5 seconds on/5 seconds off) for 5 minutes. Lysate was clarified by centrifugation at 20,000 rpm for 40 minutes at 4 °C and filtration through a 0.45 μm PVDF syringe filter. Supernatant was applied to a 5 mL HisTrap HP column (Cytiva #17524801) equilibrated with Ni-NTA Buffer A (20 mM HEPES pH 7.5, 500 mM NaCl, 5 mM β-mercaptoethanol, 10 μM ZnCl_2_, 10 mM imidazole). Column was washed with a linear gradient of 0-10% Ni-NTA Buffer B (20 mM HEPES pH 7.5, 500 mM NaCl, 5 mM β-mercaptoethanol, 10 μM ZnCl_2_, 300 mM imidazole) over 11 CV, and eluted using a linear gradient of 60-100% Ni-NTA Buffer B over 6 CV. RPA protein eluted at approximately 70-80% Ni-NTA Buffer B. Eluent was incubated with H3C protease for 1 hour at room temperature to cleave the His_6_ tag and dialyzed in Pre-Heparin Buffer (20 mM HEPES pH 7.5, 300 mM NaCl, 5 mM β-mercaptoethanol, 10 μM ZnCl_2_, 10% glycerol) for 3 hours at 4 °C. Dialyzed, cleaved protein was diluted to a final salt concentration of 150 mM NaCl using Dilution Buffer (20 mM HEPES pH 7.5, 5 mM β-mercaptoethanol, 10 μM ZnCl_2_, 10% glycerol). Sample was applied to a 5 mL HiTrap Heparin HP column (Cytiva #17040701) equilibrated with Heparin Buffer A (20 mM HEPES pH 7.5, 50 mM NaCl, 5 mM β-mercaptoethanol, 10 μM ZnCl_2_, 10% glycerol), washed with a linear gradient of 0-15% Heparin Buffer B (20 mM HEPES pH 7.5, 1 M NaCl, 5 mM β-mercaptoethanol, 10 μM ZnCl_2_, 10% glycerol) over 6 CV, 15% Heparin Buffer B over 8 CV, and eluted using a linear gradient of 15-50% Heparin Buffer B over 10 CV, followed by 100% Heparin Buffer B over 6 CV. RPA protein eluted at approximately 25% Heparin Buffer B. Eluted protein was further purified on a 24 mL Superdex 200 Increase 10/300 GL column (Cytiva #28-9909-44) equilibrated with S200 Buffer (20 mM Tris pH 7.5, 200 mM NaCl). RPA protein eluted at approximately 13 mL. Protein identity was confirmed by electrospray ionization mass spectrometry and used for experimentation within 24 hours after final elution, or aliquoted and flash frozen in liquid nitrogen.

## *In Vitro* NER Activity Assay with Purified Proteins

A plasmid substrate containing a site-specific 1,3-GTG cisplatin intrastrand crosslink was generated from p98 plasmids employing gapped-vector technology using a purified 24-mer of the platinated oligonucleotide (5’-pTCTTCTTCT**GTG**CACTCTTCTTCT). (25) The plasmid containing and the essential purified, recombinant NER proteins except XPA required to reconstitute dual-incision activity *in vitro* was incubated with either wild-type or variant XPA as described previously. (13) For each reaction, 5 nM XPC-RAD23B, 40 nM RPA, 27 nM XPG, 13.3 nM XPF-ERCC1, and 10 nM TFIIH was complemented with 20 nM WT or variant XPA. All proteins were > 95% pure. Full-length XPA and RPA were produced as described in Materials and Methods, and all remaining proteins were produced as previously described: XPC-RAD23B, (14) XPG, (15) XPF-ERCC1, (16) and TFIIH. (17) Incision reactions were conducted in repair buffer (45 mM HEPES-KOH pH 7.8, 5 mM MgCl_2_, 0.3 mM EDTA, 40 mM phosphocreatine di-Tris salt, 2 mM ATP, 1 mM DTT, 2.5 μg/μL BSA, 0.05 μg/µL creatine phosphokinase, and 70 mM NaCl) at 30 °C for indicated times. A 3’-phophorylated oligonucleotide (0.5 μL of 1 μM solution) was added for product labeling and the mixture heated at 95 °C for 5 min. The mixture was cooled to room temperature over 15 min. 1.2 μL of a Sequenase/[α^32^P]-dCTP mix (0.25 units of Sequenase and 2.5 μCi of [α32P]-dCTP per reaction) was added and incubated at 37 °C for 3 min. Then 1.2 μL of dNTP mix (100 μM of each dATP, dTTP, dGTP; 50 μM dCTP) was added to mixture and incubated for another 12 min. The reactions were stopped by adding 12 μL of loading dye (80% formamide/10 mM EDTA) and heating at 95 °C for 5 min. 6 μL of sample was loaded on 14% sequencing gel (7 M urea, 0.5x TBE) and electrophoresed at 45 W for 2.5 hrs. The reaction products were visualized using a PhosphorImager (Amersham Typhoon RGB, GE Healthcare Bio-Sciences). Two independent repetitions were performed. The NER products were quantified by ImageQuant TL and normalized to the amount of NER product formed with WT-XPA at 45 min.

## Circular Dichroism (CD) Spectroscopy

CD spectra were recorded and thermal denaturation monitored by CD for indicated XPA DBD proteins similarly as described previously, (26–28) using a Jasco J-810 spectropolarimeter and a quartz cuvette. Protein samples were used within 24 hours after purification or thawed from flash frozen aliquots, exchanged into 150 mM KH_2_PO_4_ buffer at pH 7.5, and 0.2 μm filtered. The far-UV spectrum was measured from 260 to 190 nm in increments of 0.5 nm at a rate of 50 nm/min with a response time of 2 seconds and a bandwidth of 1 nm. For each independent experiment, three spectra were averaged and smoothed for each protein and a buffer blank spectrum was subtracted.

For thermal denaturation, CD was monitored every 1.0 °C at 222 nm as the temperature was increased from 20 to 80 °C for 90 minutes, and the first derivative curve was generated for the resulting buffer blank-subtracted curve. The mean apparent T_m_ was determined by taking the average of the temperatures corresponding to the maximum values from each first derivative curve for three independent denaturation experiments.

## Microscale Thermophoresis (MST)

MST to measure DNA binding affinity was performed as described previously. (13,29) DNA binding affinity of indicated XPA DBD proteins for a 5’ 6-FAM-labeled 8 nucleotide (nt) hairpin substrate with a 4 nt overhang (5’-6-FAM-TTTTGCGGCCGCTTTTGCGGCCGC-3’) was measured using a Monolith NT.115 (NanoTemper Technologies). DNA substrate was prepared for use by diluting to working concentration of 440 nM in Substrate Buffer (6 mM Tris pH 7.5, 60 mM KCl, 0.2 mM MgCl_2_), heating at 95 °C for 5 minutes, and cooling on ice. Indicated XPA proteins were used within 24 hours after purification or thawed from flash frozen aliquots and filtered using 0.2 mm wwPTFE Pall Nanosep centrifugal filters (Pall #ODPTFE02C35). Protein samples were buffer exchanged into XPA MST Buffer (50 mM Tris-HCl pH 7.8, 150 mM NaCl, 10 mM MgCl_2_, 0.05% Tween-20, 1 mM DTT) and diluted into 16 concentrations ranging from 0.1 to 50 μM on ice. Immediately prior to measuring affinity, DNA substrate was added to each protein sample to a final concentration of 40 nM. All measurements for each independent replicate were carried out with four technical replicates for each protein concentration.

Measurements were taken at room temperature using standard capillaries (NanoTemper Technologies #MO-K022) with MST power set to low and laser excitation power set to 20%. Fraction of substrate bound at each XPA concentration was reported using values output by the MO.AffinityAnalysis software (NanoTemper Technologies). Y148D XPA DBD could not be concentrated to the final 50 μM datapoint due to limited solubility.

## Isothermal Titration Calorimetry (ITC)

ITC was performed with purified full-length XPA and RPA protein samples as described previously. (13) Protein samples were used within 24 hours after purification or thawed from flash frozen aliquots and dialyzed into ITC Buffer (20 mM Tris pH 8.0, 150 mM NaCl, 3% glycerol, 0.5 mM tris(2-carboxyethyl)phosphine (TCEP)). All buffers and protein samples were 0.2 μm filtered and degassed prior to beginning. Titrations were performed at 25 °C with 125 rpm stirring using a Affinity ITC instrument (TA Instruments), and included an initial injection of 0.5 μL of 115 μM XPA into the sample cell containing 20 μM RPA, followed by an additional 47 injections of 3 μL each. Injections were spaced over 200 – 250 second intervals. Data were analyzed using NanoAnalyze (TA Instruments) and the thermodynamic parameters and binding affinities were calculated using the average of two or three titrations fit to an independent binding model.

## Local UV Irradiation Assay

Local UV irradiation was performed similarly to as described previously. (13,30) Cells were plated onto glass coverslips (Millipore Sigma #CLS285022) coated with 1 μg/mL fibronectin (Millipore Sigma #F4759) at a density of 5.0 x 10^5^ cells per well of a 6-well plate and incubated overnight following standard culture conditions. Adherent cells were washed once with 1X DPBS (Thermo Fisher Scientific #14190-144) and the coverslip was removed and placed cell-side up onto a clean 10 cm dish covered with parafilm for UV irradiation. A polycarbonate isopore membrane filter with 5 μm pores (Millipore Sigma #TMTP04700) was pre-soaked in 1X DPBS and placed directly on top of the coverslip, and cells were irradiated with 120 J/m^2^ of UV-C. Post-irradiation, the membrane filter was gently removed and the coverslip was returned to a 6-well plate with fresh culture media and incubated following standard culture conditions for 30 minutes. Coverslips were washed once with cold 1X PBS and cells permeabilized with Hypotonic Lysis Buffer (10 mM Tris-HCl pH 8.0, 2.5 mM MgCl2, 10 mM β-glycerophosphate, 0.2 mM PMSF, 0.1 mM Na_3_VO_4_, 0.1% Igepal) for 8 minutes on ice. Coverslips were washed with Hypotonic Lysis Buffer without Igepal for 4 minutes at room temperature. Cells were fixed with 4% paraformaldehyde (Fisher Scientific #50-980-487) freshly diluted in 1X PBS (137 mM NaCl, 2.7 mM KCl, 10 mM Na_2_HPO_4_, 1.8 mM KH_2_PO_4_, pH 7.4) for 15 minutes at room temperature with gentle shaking. Permeabilized, fixed cells were incubated with 0.07 M NaOH in 1X PBS for 3 minutes at room temperature to denature DNA and then washed 5 times for 3 minutes each time with 1X PBS. Cellular localization for proteins of interest was then visualized after performing the described immunofluorescence protocol and imaging of coverslips using a Zeiss fluorescence microscope and X-Cite fluorescence lamp. The percent cells with co-localized foci was determined for each independent replicate by quantifying the number of cells with overlapping XPA and 6-4PP foci out of 100 cells with 6-4PP foci using FIJI.

## Immunofluorescence

For immunofluorescence, coverslips were blocked in 1% bovine serum albumin (BSA, Millipore Sigma #A2153) with 10% normal goat serum (Vector Laboratories #S-1000-20) in 1X PBS for one hour at room temperature. Coverslips were incubated with indicated primary antibodies in 1X PBST (1X PBS with 0.1% Tween-20) with 1% BSA and 2% normal goat serum for 2 hours at room temperature in humid conditions. Primary antibody solution was removed and coverslips were washed 3 times for 5 minutes each with 1X PBST. Coverslips were incubated with indicated secondary antibodies in 1X PBST with 1% BSA and 2% normal goat serum for 1 hour at room temperature in dark and humid conditions. Secondary antibody solution was removed and coverslips were washed 2 times for 5 minutes each with 1X PBST. Coverslips were washed for 5 minutes in 1X PBS with 300 nM 4’-6’-damino-2-phenylindole (DAPI, dihydrochloride, Millipore Sigma #D9542). Coverslips were briefly rinsed twice with 1X PBS, mounted onto microscope slides with ProLong Diamond Antifade Mountant (Thermo Fisher Scientific #P36970), and sealed with nail polish.

## Statistical Analyses

Statistical analyses were performed in GraphPad Prism (RRID:SCR_002798). Details on sample sizes, tests used, error bars, and statistical significance values are provided in the figures and figure legends.

## Data Availability

The data generated in this study are available within the article and its supplementary files. Unique reagents including plasmids generated are available upon request to the corresponding author.

## Results

### Cells stably expressing predicted NER-deficient tumor variants have increased sensitivity to UV and cisplatin

All five XPA tumor variants selected for this analysis were predicted to have a high probability of being NER-deficient, P(NER-deficient), values of F112C = 0.962, M113I = 0.75, D114Y = 0.978, R130I = 0.993, Y148D = 0.849, (11) and thus also predicted to sensitize cells to UV light and Pt-based chemotherapeutics. Moreover, each of the mutations are located in functionally relevant regions of the protein: F112, M113, and D114 are at an essential XPA-RPA interaction interface that is required for NER activity, (13,29) R130 contacts the DNA substrate, (29) and Y148 is a buried hydrophobic residue within the globular core of the XPA DNA binding domain (DBD). (31–34)

The five XPA variants were stably expressed in SV40-transformed human dermal fibroblast XP2OS cells, which lack expression of wild-type (WT) XPA protein. (12) Stable, lentivirus-mediated expression of either WT or variant XPA was confirmed by Western blot (**Supplementary Figure S1A**). As expected, the overexpression of XPA in these cells did not alter cell proliferation in any of the cell lines (**Supplementary Figure S1B**). Although the majority of variants were expressed at levels similar to WT protein, there was a ten-fold reduction in Y148D XPA protein compared to WT protein. Control experiments showed this was not due to a difference in lentiviral transduction efficiency or issues with mRNA expression (**Supplementary Figure S1C**). Protein degradation via the Ubiquitin Proteasome System (UPS) is a common mechanism for regulating protein levels and as a means to rid the cell of misfolded or destabilized proteins. (35,36) To test whether the Y148D mutation leads to more ready degradation via the UPS, cells stably expressing WT or Y148D XPA as well as mock XPA-null cells were treated with the UPS inhibitor bortezomib (37,38) and levels of XPA were assessed by Western blot. UPS inhibition led to a partial rescue of Y148D protein levels (**Supplementary Figure S1D-E**). These data indicate that the lower level of Y148D is mediated at least in part by degradation by the UPS, and that this must be accounted for in our subsequent analyses.

In order to determine whether these XPA variants cause NER defects in the context of damage to native chromatin, cells were first tested for hypersensitivity to UV light using a clonogenic cell survival assay. Compared to cells expressing WT XPA, cells expressing F112C or R130I XPA did not exhibit increased sensitivity to UV. In contrast, cells expressing M113I, D114Y, and Y148D XPA showed a statistically significant increase in sensitivity to UV irradiation (IC_50_ [95% confidence interval (CI)] = 5 J/m^2^ [4 – 6 J/m^2^], 2 J/m^2^ [2 – 3 J/m^2^], 3 J/m^2^ [2 – 4 J/m^2^], 2 J/m^2^ [2 – 3 J/m^2^], for WT, M113I, D114Y, and Y148D, respectively) (**Figure 1A**, **Supplementary Figure S2**). Reassuringly, this data agreed with our previous findings using a UV-damaged reporter plasmid in a host-cell reactivation assay (FM-HCR), (11) which showed these variants have reduced capacity for the repair of UV lesions.

**Figure 1.**
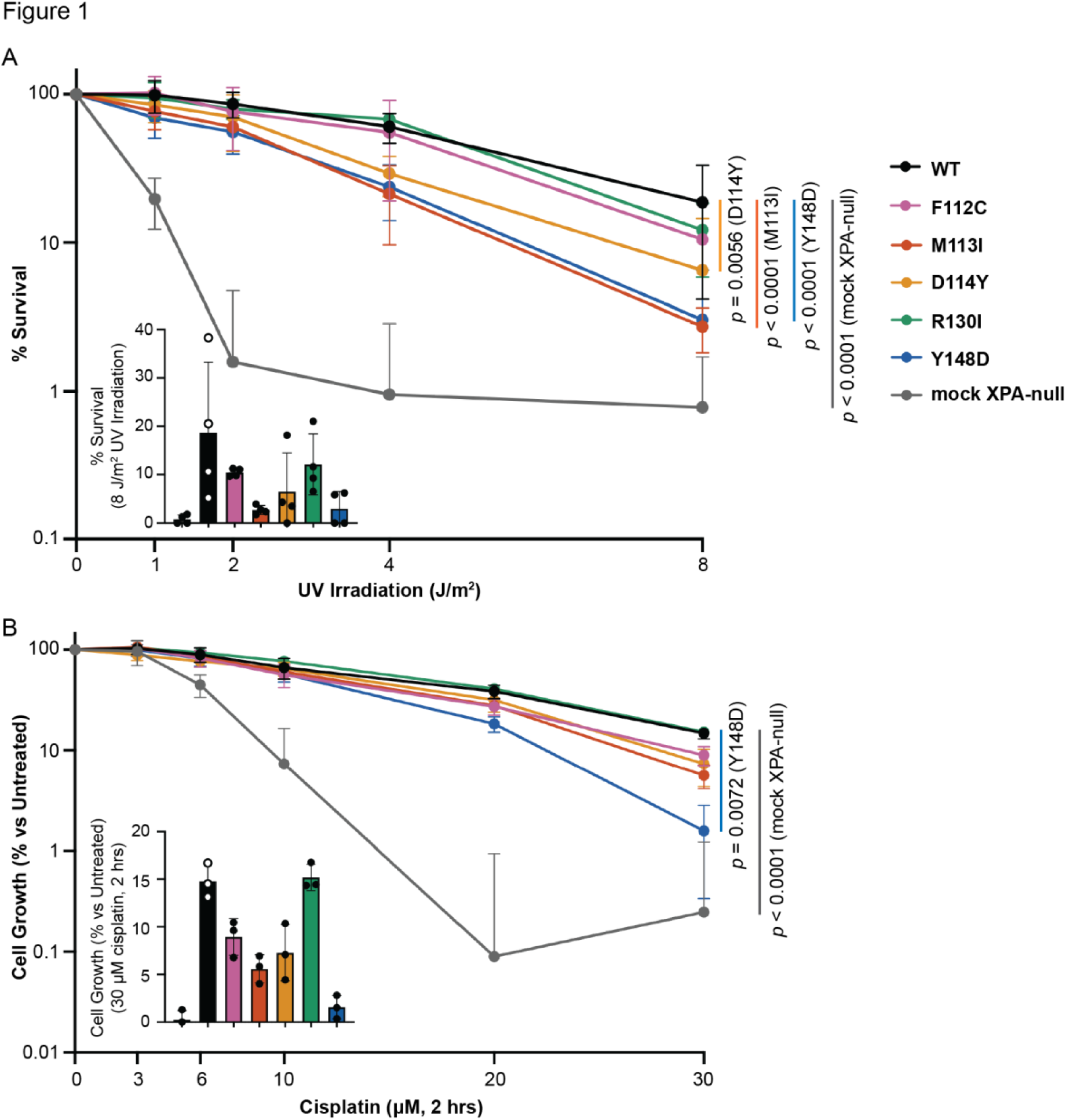
**Cells stably expressing NER-deficient XPA tumor variants have increased sensitivity to UV and cisplatin. *A***, Clonogenic cell survival assay in stable XP2OS cell lines after exposure to UV at indicated doses and growth for 14 days (n = 4). Colonies were stained with 0.5% crystal violet and those greater than approximately 50 cells were counted using Fiji. Percent survival was determined for each cell line relative to untreated. IC_50_ values [95% confidence intervals (CI)] determined using lines of best fit from nonlinear regression analysis in GraphPad Prism ([inhibitor] vs. response – variable slope (four parameters equation): 0.6 J/m^2^ [0.2 – 0.8 J/m^2^] (mock XPA-null), 5 J/m^2^ [4 – 6 J/m^2^] (WT), 4 J/m^2^ [3 – 5 J/m^2^] (F112C), 2 J/m^2^ [2 – 3 J/m^2^] (M113I), 3 J/m^2^ [2 – 4 J/m^2^] (D114Y), 5 J/m^2^ [4 – 6 J/m^2^] (R130I) and 2 J/m^2^ [2 – 3 J/m^2^] (Y148D). EC50 shift, X is concentration equation in GraphPad Prism used to compare the IC_50_ from each line of best fit to that of the WT control, indicating comparisons to WT with *p* < 0.05. Inset showing mean percent survival for each cell line at 8 J/m^2^ UV irradiation. ***B***, SRB assay in stable XP2OS cell lines after exposure to cisplatin for two hours at indicated doses and growth for five days (n = 3). Cells were stained with 0.4% SRB and absorbance at 490 nm measured. Percent untreated cell growth was determined for each cell line relative to untreated. IC_50_ values [95% CI] determined using lines of best fit from nonlinear regression analysis in GraphPad Prism ([inhibitor] vs. response – variable slope (four parameters equation): 6 μM [5 – 7 μM] (mock XPA-null), 15 μM [12 – 18 μM] (WT), 12 μM [11 – 15 μM] (F112C), 13 μM [11 – 15 μM] (M113I), 13 μM [11 – 16 μM] (D114Y), 17 μM [15 – 20 μM] (R130I) and 11 μM [10 – 14 μM] (Y148D). EC50 shift, X is concentration equation used to compare the IC_50_ of each line of best fit to that of the WT control, indicating comparisons to WT with *p* < 0.05. Legend in panel (***A***). Inset showing mean cell growth for each cell line at 30 μM cisplatin.

A similar analysis of cisplatin hypersensitivity of two sets of independently generated stable cell lines was performed using a sulforhodamine B assay. These experiments revealed that only cells expressing the Y148D variant XPA had a statistically significant increase in cisplatin sensitivity compared to WT XPA-expressing cells: cell line set 1 (IC_50_ [95% CI] = 11 μM [10 – 14 μM] for Y148D XPA versus 15 μM [12 – 18 μM] for WT XPA) (**Figure 1B**), and cell line set 2 (IC_50_ [95% CI] = 14 μM [11 – 15 μM] for Y148D XPA versus 18 μM [15 – 20 μM] for WT XPA) (**Supplementary Figure S3A-B**). Some cisplatin lesions can be repaired by non-NER pathways including double-strand break (DSB) repair and interstrand crosslink (ICL) repair, (39) which may explain the differences observed between repair of UV and cisplatin lesions in the cellular assays. Regardless, our results suggest that expression of the Y148D XPA tumor variant in cells leads to decreased efficiency in the repair of both UV and cisplatin lesions.

### XPA tumor variants are defective in dual-incision of a NER-specific cisplatin lesion in vitro

To further address the difference between the UV and cisplatin sensitivity and obtain deeper insights into the functionality of XPA tumor variants, an *in vitro* repair assay with a NER-specific 1,3-GTG cisplatin intrastrand crosslink lesion (40–43) was performed. Purified recombinant NER proteins RPA, XPC-RAD23B, XPG, XPF-ERCC1, and TFIIH were complemented with equal concentrations of WT or variant XPA protein and the appearance of NER excision products was monitored in a time-dependent manner. Importantly, as opposed to the assays performed in stable cell lines, this *in vitro* reconstitution also allowed direct comparison of the effect of different variants on NER that was independent of native cellular concentration. This was particularly important for the comparative analysis of Y148D. Consistent with the decreased levels of Y148D observed in cells (**Supplementary Figures S1A** and **S3A**), we note that the recombinant protein was expressed primarily in the insoluble fraction and required unique approaches to obtain sufficient quantities of protein for use in this assay.

Reconstitution of the NER reaction *in vitro* without XPA did not lead to excision products for the damaged substrate, whereas addition of WT, F112C, or R130I XPA led to robust accumulation of excision products over time (**Figure 2A-B**). These results agreed with the lack of hypersensitivity to UV or cisplatin damage in cells expressing WT, F112C, or R130I XPA. Addition of either M113I or D114Y XPA to the reaction led to a notable but not statistically significant decrease of ∼25% of excision product accumulation compared to WT XPA (**Figure 2A-B**). We note that this effect is somewhat reduced relative to that observed in the survival assays (**Figure 1A**), which we attribute to the limitations of an *in vitro* reconstitution biochemical assay relative to the cellular context. This finding suggests that the M113I and D114Y mutations introduce a functional defect in XPA during NER of cisplatin lesions, although this was masked in the cellular assays. In the case of the Y148D variant, a much larger ∼50% decrease of excision product accumulation by the final 45-minute timepoint (**Figure 2A-B**) was observed, independent of the decrease in protein seen in cells. These results suggest that the Y148D XPA variant leads to a functional protein defect in addition to increased cellular degradation. To obtain deeper insights, we set out to determine the mechanisms of NER dysfunction for these tumor variants by searching for evidence of perturbation of protein structure and stability, decreased DNA binding affinity, loss of protein-protein interaction, and disruption of recruitment to sites of damage.

**Figure 2.**
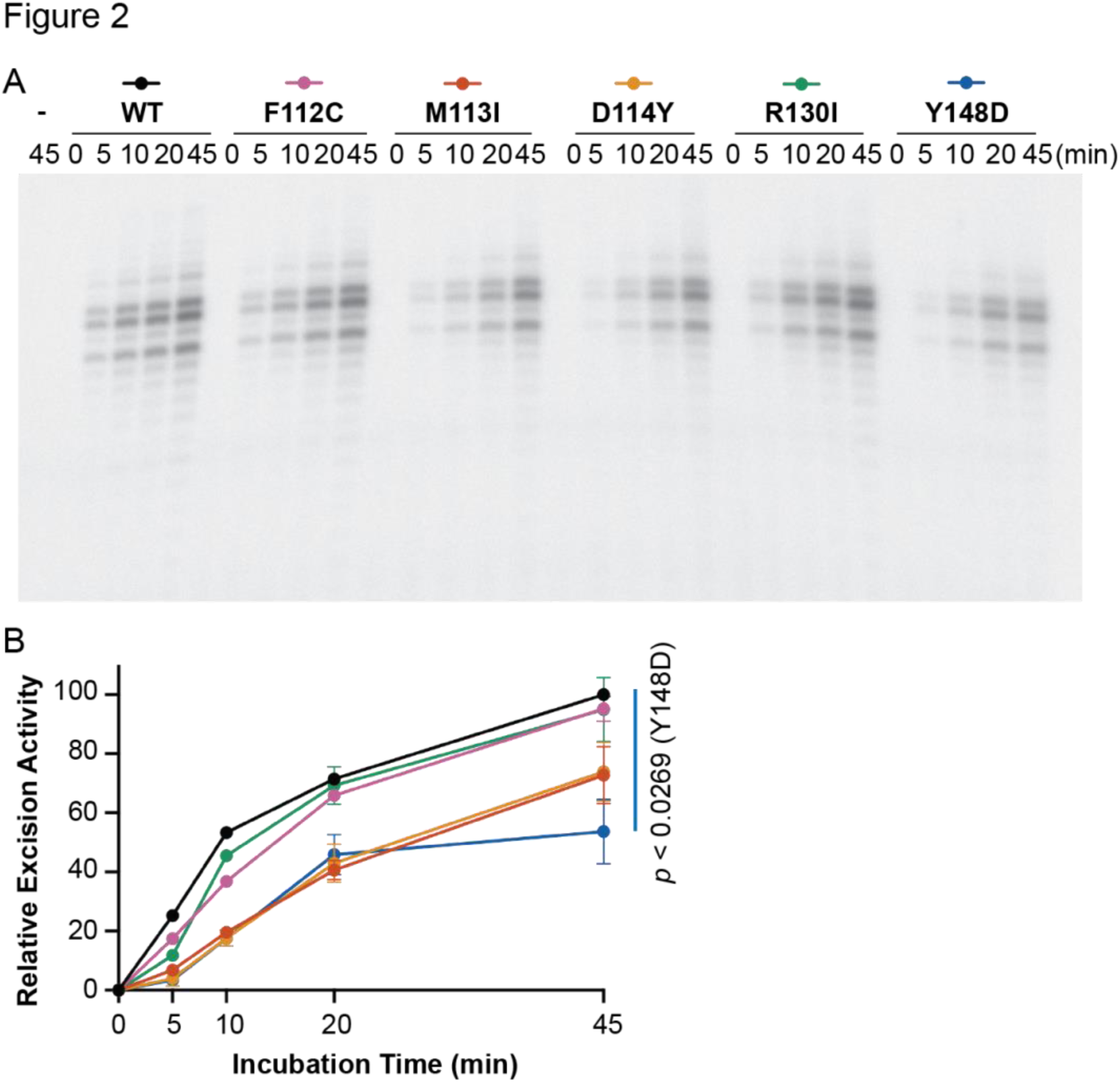
**XPA tumor variants are defective in dual-incision of a NER-specific cisplatin lesion *in vitro*. *A***, *In vitro* NER activity of purified NER proteins RPA (40 nM), XPC-RAD23B (5 nM), XPG (27 nM), XPF-ERCC1 (13 nM), and TFIIH (10 nM) complemented with WT or variant XPA (20 nM) on a plasmid containing a 1,3-GTG cisplatin intrastrand crosslink (n = 2). Gel shows excision products. ***B***, Excised band intensities for the reconstituted complexes containing each different XPA protein, as indicated by the legend in ***A***. Relative excision activity was determined as percentage of the WT XPA excision activity at 45 minutes. Mean relative excision activity values at 45 minutes were statistically compared for each variant to WT using two-tailed unpaired t-tests, and comparisons to WT with *p* < 0.05 are indicated.

### The Y148D mutation destabilizes XPA

A fundamental step in defining the molecular mechanisms of dysfunction for a variant is to determine if the mutation has any effect on the structural stability of the protein. To this end, we turned to circular dichroism (CD) spectroscopy to characterize the distribution and stability of secondary structural elements for the XPA variants. The previously well-characterized XPA DNA binding domain (DBD, XPA_98-239_) was used for these experiments because it is the only part of the XPA that folds into a globular domain with regular secondary structure and all the variants investigated are within this domain. XPA DBD consists primarily of α-helical elements, (33,44–46) a characteristic that is well reflected in the CD data (**Supplementary Figure S4**), which show that all five variants retain the structure of the WT protein. The effect of the mutations on the thermal stability of XPA DBD was then assayed by measuring the thermal denaturation midpoint (T_m_) derived from the temperature dependence of the CD spectra.

Compared to WT, the M113I and R130I mutations had no effect on T_m_ (59 ± 0.6 °C, 59 ± 0.6 °C and 58 ± 1.0 °C for WT, M113I and R130I, respectively), whereas F112C, and D114Y had statistically significant but very modest decreases in T_m_ (53 ± 0.0 °C and 57 ± 0.0 °C for F112C and D114Y, respectively) (**Figure 3**). The T_m_ decrease of only a few degrees for either F112C or D114Y is highly unlikely to cause significant defects in the context of NER function. In contrast, the Y148D variant was dramatically destabilized (T_m_ 38 ± 2.1 °C) to the extent that the protein will have a significant population of unfolded protein in cells, consistent with our observations of lower levels and degradation by the UPS in cells.

**Figure 3.**
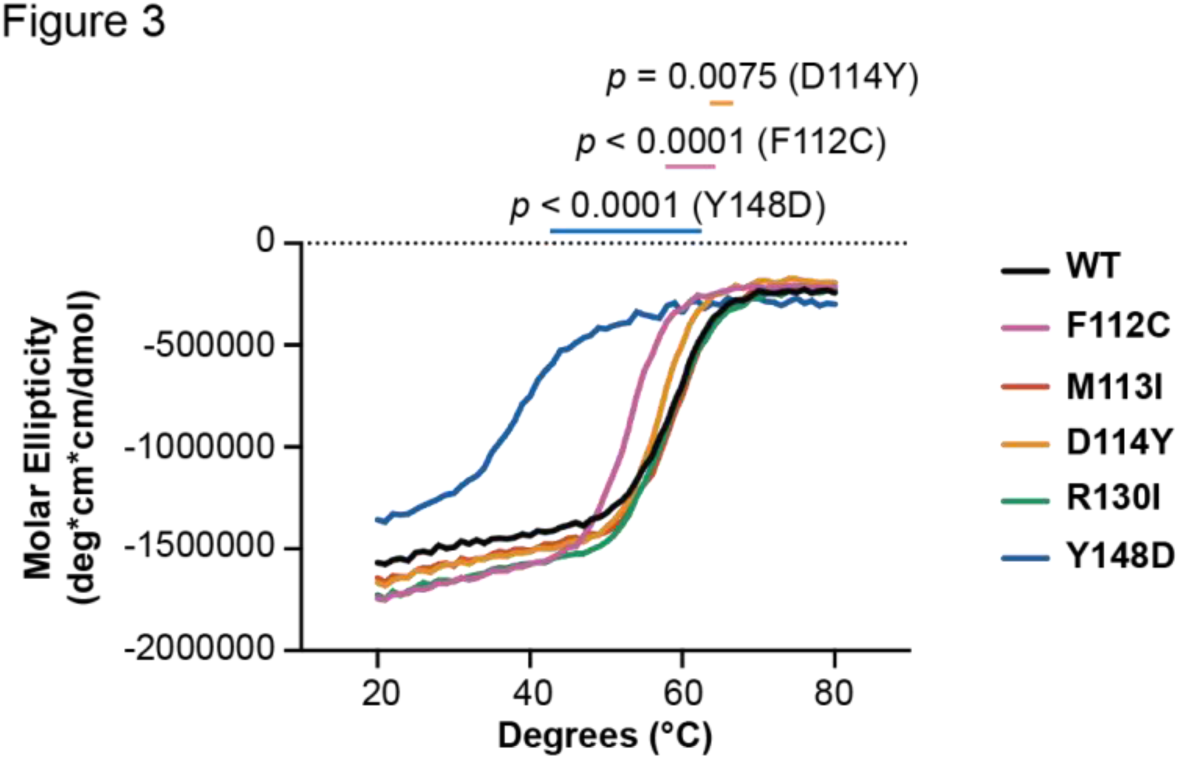
The Y148D variant destabilizes XPA. Circular dichroism of purified recombinant WT or variant XPA DBD protein measured at 222 nm over increasing temperature (n = 3). Each curve represents the average of three measurements. Ellipticities for the buffer and cuvette alone were subtracted from each measurement. Mean apparent T_m_ and standard deviation determined for each variant by recording the temperature at which the steepest slope is observed (the temperature at the maximum recorded value of the corresponding first derivative curve): 59 ± 0.6 °C (WT), 53 ± 0.0 °C (F112C), 59 ± 0.6 °C (M113I), 57 ± 0.0 °C (D114Y), 58 ± 1.0 °C (R130I) and 38 ± 2.1 °C (Y148D). Mean apparent T_m_ values were statistically compared for each variant to WT using two-tailed unpaired t-tests, and comparisons to WT with *p* < 0.05 are indicated.

### XPA tumor variants have disrupted DNA binding affinity

The binding of DNA by XPA is central to its function as a scaffold in NER, so the DNA binding affinity was measured for each XPA variant and compared to WT protein. Microscale thermophoresis (MST) with XPA DBD and a fluorescently labeled NER junction mimic substrate was used for these measurements. (29) The DNA binding affinity observed for WT DBD was comparable to that reported previously (K_d_ = 7.7 ± 3.7 μM). (13,29,47) Among the five XPA variants, there was no effect on DNA binding affinity for F112C, M113I and D114Y (**Figure 4**), although this was perhaps expected because these residues do not contact DNA and instead are located within the RPA interaction interface on XPA. For R130I, a residue known to contact the DNA substrate, (29) we observed a statistically significant but very modest reduction in affinity (K_d_ = 38 ± 22 μM). The small size of the effect reflects the limited contribution of any single residue within the large DNA binding surface on XPA, and is consistent with the lack of hypersensitivity to the UV and cisplatin. In contrast to the other variants, Y148D did not have any measurable affinity for DNA (**Figure 4**), suggesting the observed NER deficiency for this variant arises from a scaffolding defect in NER.

**Figure 4.**
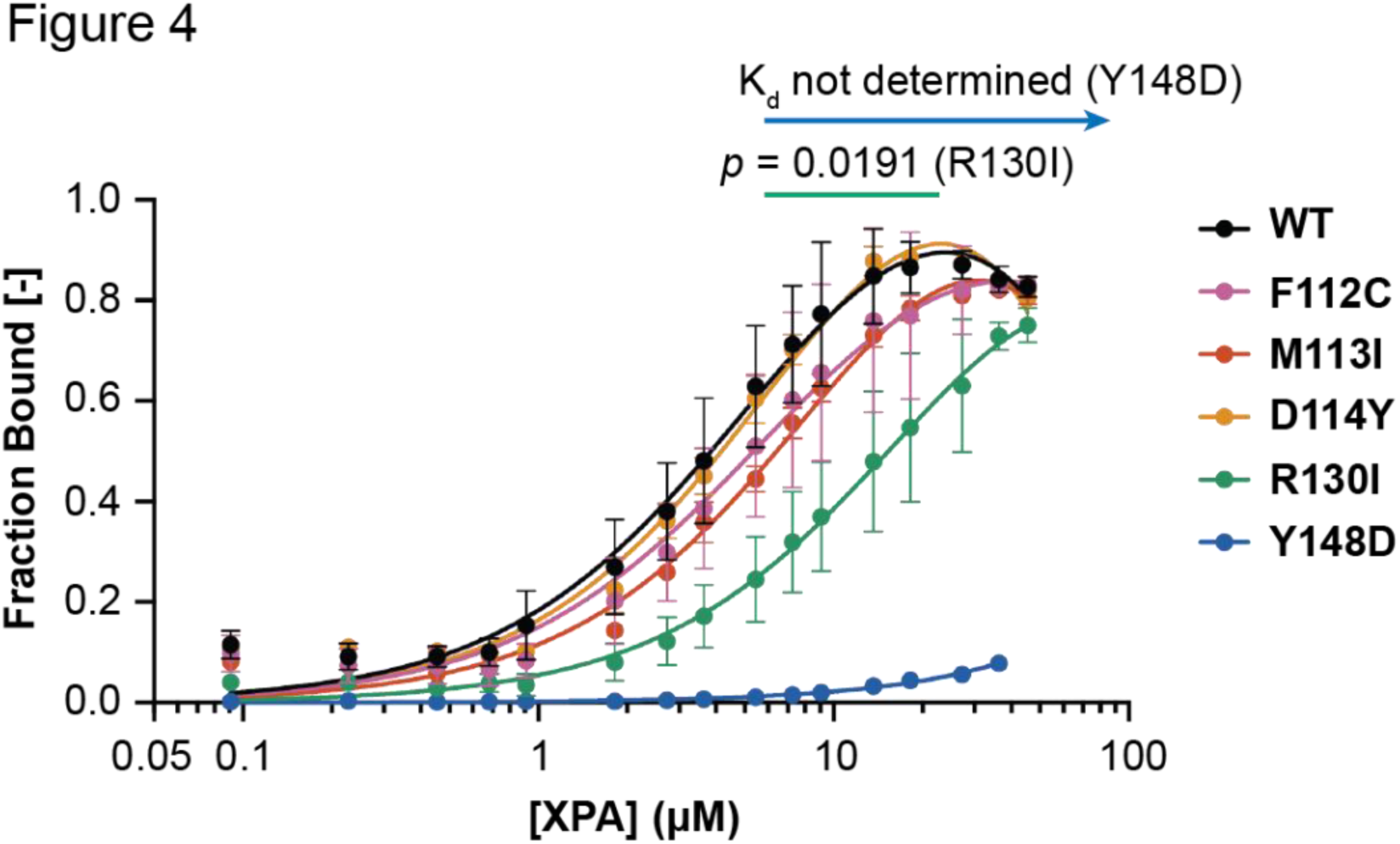
XPA tumor variants have disrupted DNA binding affinity. Microscale thermophoresis using a 5’ 6-FAM labeled hairpin DNA substrate with 8 nucleotides (nt) of dsDNA and 4 nt of ssDNA overhang and indicated WT or variant purified recombinant XPA DBD (residues 98-239) protein performed at 20 °C (n = 5 for WT, n = 3 for all variants). K_d_ values for the DNA substrate in each replicate were determined using lines of best fit from nonlinear regression analysis in GraphPad Prism (one site – total binding model). Error bars indicate the standard deviation. The mean and standard deviation K_d_ values were 7.7 ± 3.7 μM (WT), 9.2 ± 2.2 μM (F112C), 12 ± 1.9 μM (M113I), 8.8 ± 1.7 μM (D114Y), 38 ± 22 μM (R130I) and not determined (Y148D). Mean K_d_ values were statistically compared for each variant to WT using two-tailed unpaired t-tests, and comparisons to WT with *p* < 0.05 are indicated.

### NER-deficient variants M113I and D114Y do not significantly disrupt interaction with RPA

We have shown previously that NER function requires interaction between XPA and RPA and that mutations within the RPA interaction interface of XPA can inhibit NER. (13,29) In the effort to discern the mechanistic basis for the NER deficiencies observed for three of our variants (M113I, D114Y and Y148D), we set out to apply our isothermal titration calorimetry (ITC) approach to test the interaction of XPA with RPA. Attempts to apply this assay to Y148D were stymied by its limited solubility, which prevented concentrating the protein to a sufficiently high level for ITC analysis. The K_d_ values for M113I (5.3 ± 1.1 μM) and D114Y (4.5 ± 4.9 μM) were within the experimental error of the value for WT XPA (2.8 ± 1.7 μM) (**Figure 5**, **Table 1**). These results are consistent with our previous findings, which showed that single mutations within the interaction interface of the XPA DBD are insufficient to disrupt the XPA-RPA scaffold. (13,29)

**Figure 5.**
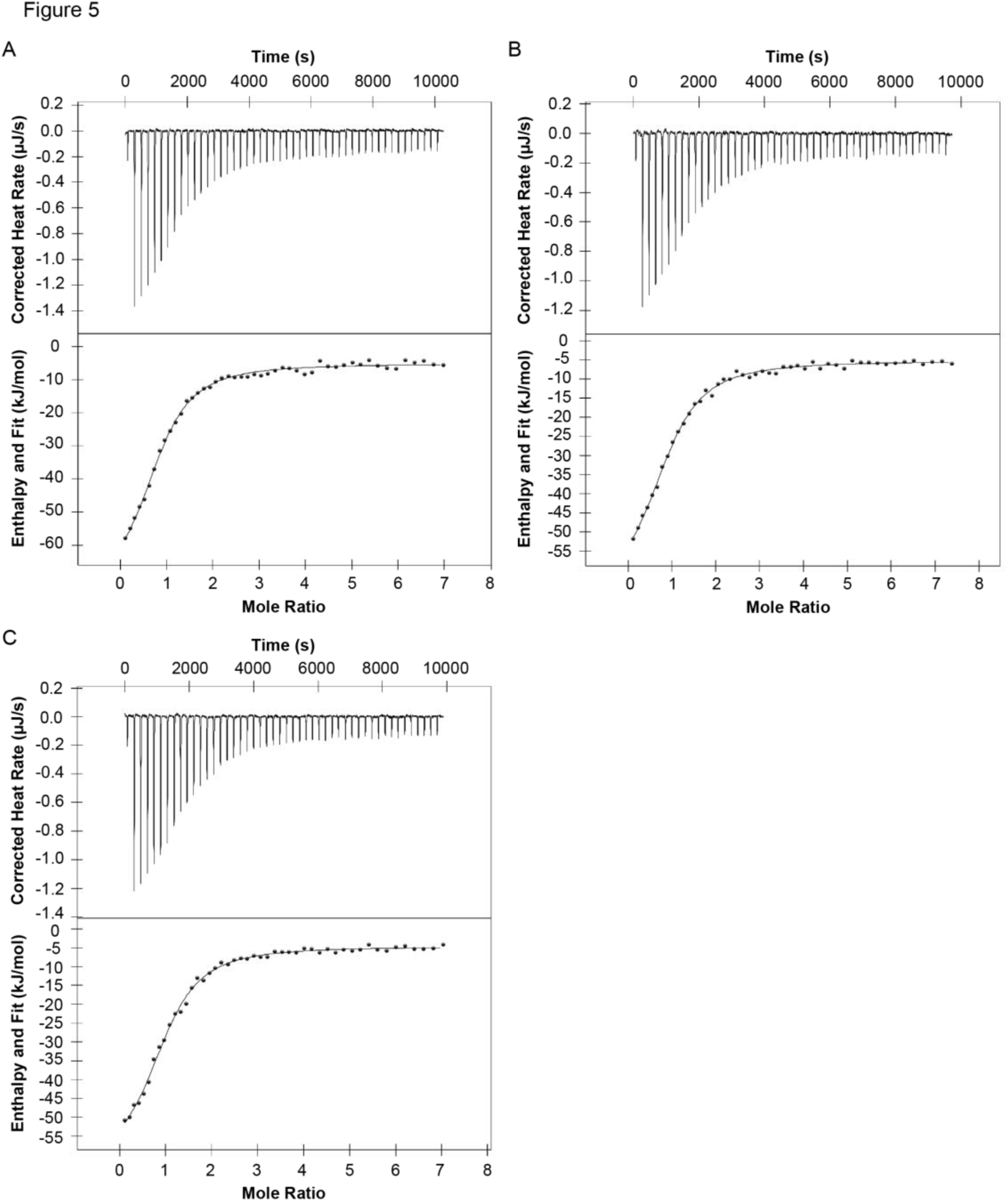
Single NER-deficient XPA variants M113I and D114Y do not significantly disrupt the essential scaffolding interaction with RPA. Isothermal titration calorimetry of purified recombinant full-length WT XPA (n = 2, *A*), M113I XPA (n = 3, *B*), and D114Y XPA (n = 2, *C*) and RPA. Representative thermograms showing the raw heat release (upper) and integrated heat release (lower) for each. The first injection of 0.5 μL was removed from each titration for analysis. The mean and standard deviation K_d_ values were 2.8 ± 1.7 μM (WT), 5.3 ± 1.1 μM (M113I), and 4.5 ± 4.9 μM (D114Y). Mean K_d_ values were statistically compared for each variant to WT using two-tailed unpaired t-tests: M113I *p* = 0.2118 and D114Y *p* = 0.4161.

**Table 1.** Thermodynamic parameters for the binding of XPA to RPA.

| XPA protein | $K_d$ (μM) | $n$ | $\Delta H$ (kJ/mol) | $-T\Delta S$ (kJ/mol) |
| --- | --- | --- | --- | --- |
| WT | $2.8 \pm 1.7$ | $1.0 \pm 0.1$ | $-56 \pm 12$ | $23.2 \pm 13.7$ |
| M113I | $5.3 \pm 1.1$ | $1.2 \pm 0.2$ | $-57 \pm 3.4$ | $26.4 \pm 3.6$ |
| D114Y | $4.5 \pm 4.9$ | $1.0 \pm 0.0$ | $-56 \pm 1.2$ | $25.5 \pm 0.9$ |

### Recruitment to sites of UV damage is hindered for Y148D XPA tumor variant

As a NER scaffold protein, XPA interacts with many other NER factors as well as RPA and the DNA near a lesion, all of which are required to assemble the catalytically competent incision complex. (48) Therefore, recruitment of XPA to the damaged site is integral to successful repair. The hypersensitivity of cells expressing M113I, D114Y, or Y148D XPA to UV and cisplatin suggests a potential recruitment defect for these variants. To assess the recruitment of each XPA protein to sites of UV damage in cells, we used local UV irradiation and immunofluorescence to detect localization of each of the five XPA variants to pyrimidine-pyrimidone (6–4) photoproducts (6-4PPs) that are classically repaired by NER. (49) Among the five variants tested, only cells expressing Y148D XPA had a statistically significant decrease in

XPA recruitment to 6-4PP damage sites compared to cells expressing WT XPA (**Figure 6A-B**, **Supplementary Figure S5**). The observation of a partial loss of recruitment for Y148D further confirms that a protein scaffolding defect in cells contributes to the NER deficiency for this variant.

**Figure 6.**
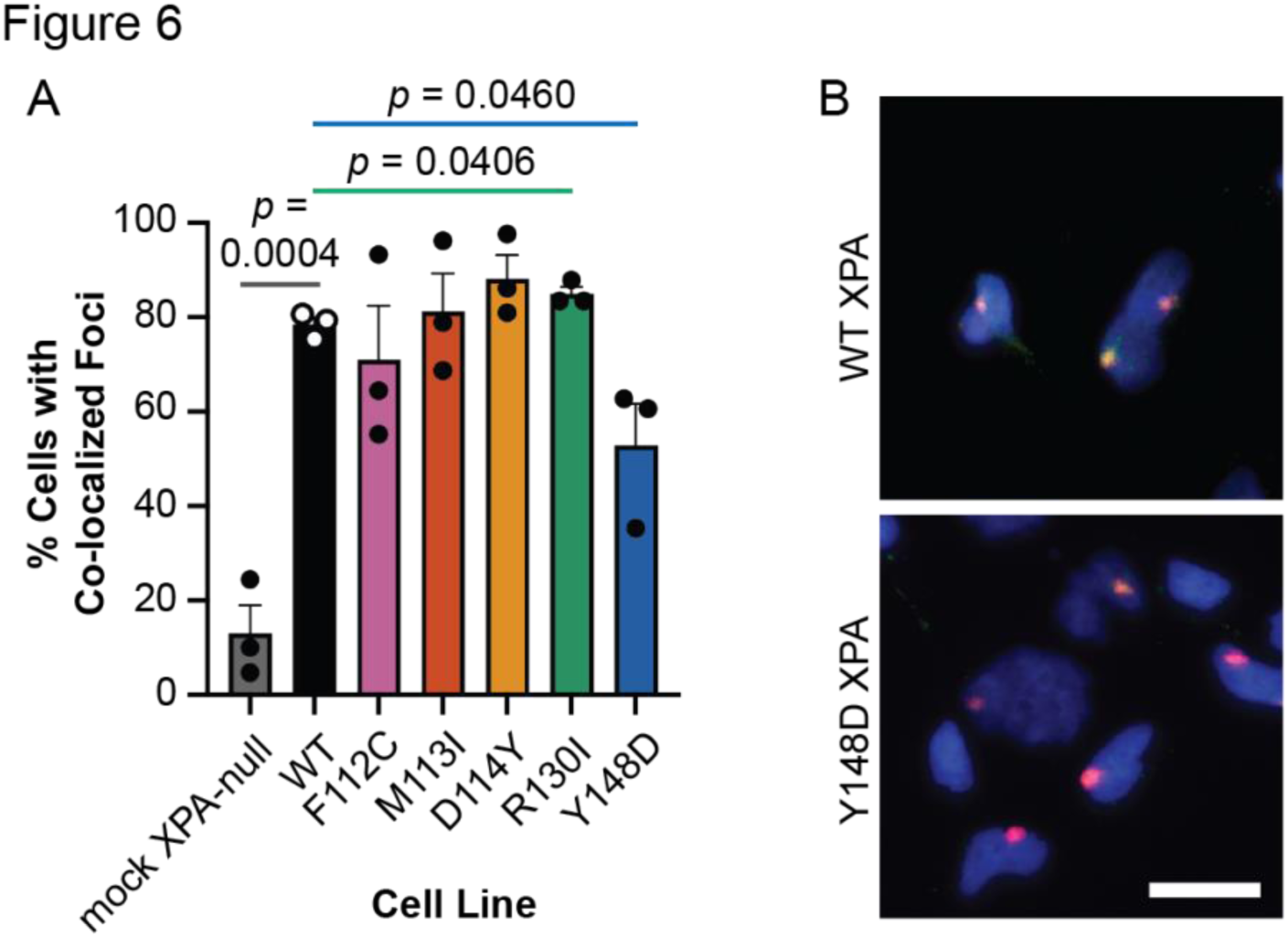
**Recruitment to sites of UV damage is hindered for Y148D XPA tumor variant. *A***, Local UV irradiation and immunofluorescence assay in stable XP2OS cell lines (n = 3). Cells were exposed to 120 J/m^2^ UV irradiation through polycarbonate isopore membranes with 5 μm pores and XPA and 6-4PP damage foci were detected using immunofluorescence after 30 minutes incubation. XPA co-localization in at least 100 cells with 6-4PP foci was quantified per cell line using FIJI. Mean percent cells with co-localized foci were statistically compared in each variant-expressing cell line to WT using two-tailed unpaired t-tests, and comparisons to WT with *p* < 0.05 are indicated. ***B***, Representative merged images of local UV irradiation and immunofluorescence assay in stable XP2OS cell lines from ***A***. DAPI in blue, 6-4PP in red, XPA in green, strong co-localization in yellow. Scale bar 20 μm.

## Discussion

The observed correlation between defects in NER genes *ERCC1* and *ERCC2* and improved outcomes for patients treated with standard-of-care Pt agents suggests that information on the status of NER genes in tumors can be used to direct patient treatments. In this study, the novel tumor variants F112C, M113I, D114Y, R130I, and Y148D were selected to test the correlation between sensitivity to UV and cisplatin induced DNA damage and elucidate mechanisms of variant dysfunction. These variants were identified by our previous work on predicting NER-deficient variants in XPA using machine learning. (11)

The NER activity of each variant was first assessed in cells with UV lesions on native chromatin (**Figure 1A**) rather than the FM-HCR reporter plasmid used in our previous study. These analyses confirmed the previous FM-HCR results that only three of the five variants of interest displayed mild (M113I and D114Y) to moderate (Y148D) sensitivity to UV in cells. Due to the preliminary nature of the machine learning predictions that identified all five variants as NER-deficient, the failure to accurately predict the F112C and R130I variants as NER-*pro*ficient was not surprising. Interestingly, only cells expressing the Y148D variant with moderate sensitivity to UV were also more sensitive to cisplatin (**Figure 1B**). This lack of agreement was somewhat unexpected based on analyses of ERCC2 variants showing near perfect correlation between UV and cisplatin sensitivity. (6) However, cisplatin lesions are known to be repaired by additional pathways in cells including DSB repair and ICL repair (39) or overcome through translesion synthesis (TLS). (50) Compensation by other pathways may have masked the mild NER-specific sensitivities to cisplatin in cells, a hypothesis supported by the *in vitro* dual-incision assay, which revealed reproducible but not statistically significant reductions in excision of a NER-specific cisplatin lesion for the M113I and D114Y variants (**Figure 2**). A similar phenomenon has been observed in ERCC1-XPA interaction mutants, where disruption of interaction with ERCC1 dramatically sensitizes cells to UV, but not cisplatin.(20) Although further investigation is necessary to better understand this discrepancy, including analysis of variants with more severe UV sensitivity as well as assays performed in cells deficient for DSB repair, ICL repair, or TLS, these findings demonstrate that XPA tumor variants such as Y148D can predictably sensitize cells to cisplatin.

To better understand how the selected XPA tumor variants led to defective NER and increased cellular sensitivity to UV or cisplatin exposure, we investigated the potential mechanisms for dysfunction of each variant. While all five of the tumor variants studied here are located in the XPA DBD and were predicted to have defective NER, increased sensitivity to UV or cisplatin was only observed for three of the variants. This highlights the importance of performing functional analyses to better understand and overcome the limitations of the initial predictive machine learning framework. Moreover, our studies to identify mechanisms of dysfunction for the remaining variants yielded important insights into essential aspects of XPA function during NER. For example, the M113I and D114Y variants, which are positioned at the XPA interaction interface with RPA, exhibited only a mild increase in UV and cisplatin sensitivity but no detectable loss in RPA binding affinity despite their location within the RPA70AB interaction interface. Both of these observations are consistent with our previous work on the interaction between the XPA DBD and RPA70AB, which revealed that single-site mutations have only a mild effect on NER activity and that multiple mutations are needed to completely abrogate this interaction. (13,29) The results obtained for the Y148D variant were more striking. Y148D was the only variant assayed that increased cellular sensitivity to both UV and cisplatin, and a severe protein stability defect was identified (**Figure 3**), as well as disrupted DNA binding (**Figure 4**) and hindered protein recruitment to sites of UV damage in cells (**Figure 6**). Considering that the Y148D variant resulted in a change from a buried hydrophobic tyrosine residue to a negatively charged aspartic acid, the resulting loss of stability reflected in the 20 °C reduction in Tm to 38 °C is not surprising. For the isolated protein this thermodynamic instability translates to approximately half of the protein being unfolded in a living cell. The Y148D gene was expressed in cells at the same level as the WT protein, but less protein was observed in cells due at least in part to turnover by the UPS, which is consistent with the protein folding defect of the variant. The DNA binding affinity measurements performed at 20 °C, a temperature much lower than the T_m_ of Y148D where the equilibrium population of unfolded protein will be substantially lower, suggest that the loss of DNA binding affinity observed is a distinct phenotype and not solely the result of protein destabilization. This conclusion is also supported by the *in vitro* incision analyses comparing equal amounts of different XPA WT and variant proteins, which indicate that the hypersensitivity phenotype is not solely due to the decreased levels of Y148D XPA protein in cells.

The analyses performed did not exhaustively assay all possible mechanisms of dysfunction. Key areas of additional study will include analyzing the rate of repair in cells expressing these variants as well as assessing the downstream recruitment of additional NER factors to sites of damage and the XPA-RPA scaffold. Nevertheless, our results provide a sound basis for the observed defects in the Y148D variant cellular hypersensitivity to UV and cisplatin. In particular, protein destabilization and degradation were identified as key mechanisms of the Y148D variant dysfunction. These insights were obtained from combining both cell-based assays and biophysical and structural analyses; neither strategy alone can provide a full explanation of variant effect on the cell. Specifically, when considering the non-enzymatic scaffolding role that XPA plays during NER, these results emphasize the importance of including protein stability metrics in future variant effect prediction efforts. More broadly, these data suggest that the predictive power of the preexisting machine learning framework can be improved by ensuring that the input training data includes metrics that reflect deleterious mechanisms known to reduce NER and sensitize cells to cisplatin as well as UV. Combined with further exploration of sensitivity to other commonly used Pt drugs such as carboplatin and oxaliplatin (51) and characterization of variants with more severe NER defects, these studies have the potential to dramatically improve XPA variant effect prediction.

Taken together, the characterization of selected XPA tumor variants predicted to sensitize cells to cisplatin revealed that XPA protein stability and scaffold function are essential for NER and that variant protein dysfunction or loss can sensitize cells to cisplatin. These findings suggest that deleterious XPA tumor variants should be considered when predicting patient response to Pt-based chemotherapeutics, in addition to those in other NER genes such as *ERCC2* or genes in other DNA repair pathways. The ability to accurately identify such variants and determine common mechanisms of dysfunction, such as destabilization and cellular degradation of XPA, remains essential for precision medicine approaches. Finally, these findings provide the foundation to test whether inclusion of mechanistic insights, such as measures of protein stability for XPA variant interpretation, significantly improve the accuracy of machine learning predictions.

## Acknowledgements

We acknowledge Anais Naretto for her assistance handling the platinum reagents. Some diagrams were created with a licensed version of BioRender.com.

## Author Contributions

Conceptualization: WJC, AMB

Methodology: AMB, KSG, MK, HSK, WJC

Software: AMB

Validation: AMB, KSG

Formal Analysis: AMB, KSG, MK, HSK, WJC

Resources: ODS, AMB

Investigation: AMB, KSG, MK, HSK, CRT, AD, AN, PDN, JP

Data Curation: N/A

Writing – Original Draft: AMB, WJC

Writing – Review and Editing: AMB, KSG, MK, HSK, ODS, WJC

Visualization: AMB, KSG, MK, HSK

Supervision: ODS, WJC

Project Administration: N/A

Funding Acquisition: WJC, ODS, AMB

## Notes

### Competing Interest Statement

The authors have declared no competing interest.

