## Supplementary Information for "XPA tumor variants lead to defects in NER that sensitize cells to cisplatin"

### Contents

Supplementary Figure S1. Stable cell lines to probe the impact of XPA tumor variants on NER and genotoxin sensitivity.
Supplementary Figure S2. Representative images of UV clonogenic survival assay in stable XP2OS cells.
Supplementary Figure S3. A second set of cells stably expressing NER-deficient XPA tumor variants has increased sensitivity to cisplatin.
Supplementary Figure S4. Global secondary structure of purified XPA variant proteins.
Supplementary Figure S5. Representative images of XPA variant co-localization with UV damage in stable XP2OS cells.
Supplementary Table 1. Mutagenesis primers to generate XPA variants in expression vectors.

Supplementary Figure S1

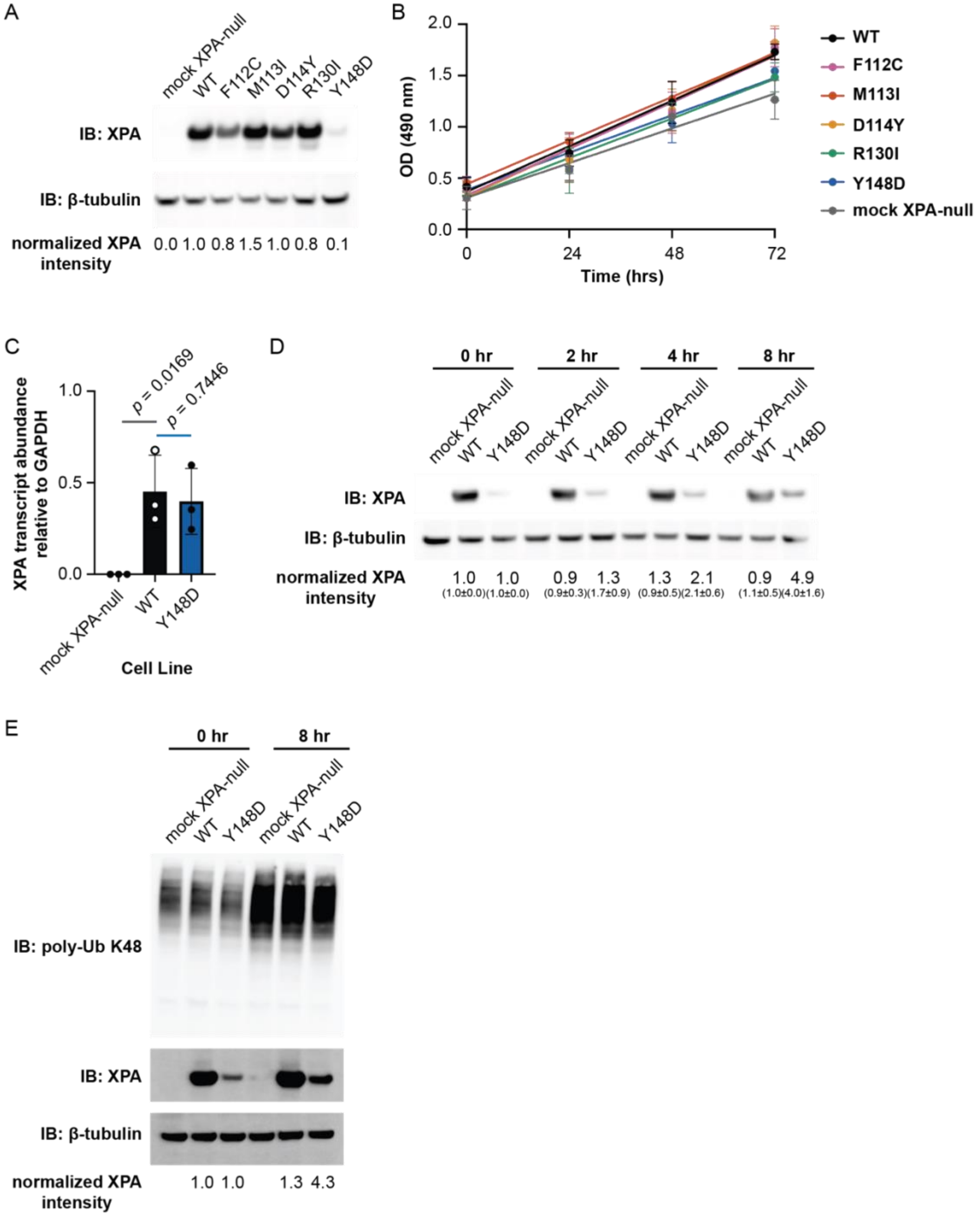


**Supplementary Figure S1. Stable cell lines to probe the impact of XPA tumor variants on NER and genotoxin sensitivity.** ***A***, Western blot analysis of stable mock XPA-null, wild-type (WT) or variant XPA overexpression levels in XP patient-derived dermal fibroblast XP2OS cells transduced with lentivirus. β-tubulin included as loading control. Band intensities were calculated using FIJI. Normalized XPA intensity determined by first normalizing the values for each XPA lane to the corresponding β-tubulin signal and then back to that for WT XPA. ***B***, SRB assay to compare cell proliferation rates of indicated stable XP2OS cell lines (n=3). Lines of best fit determined by performing a linear fit using GraphPad Prism nonlinear regression analysis, with comparison of fit test to determine whether the best-fit values of slope differ between cell lines ($p = 0.2151$). ***C***, RT-qPCR to measure *XPA* transcript abundance normalized to *GAPDH* in indicated stable XP2OS cell lines (n=3). Mean values were statistically compared to WT with a two-tailed unpaired t-test. ***D***, Representative Western blot images from analysis of stable XP2OS cells expressing mock XPA-null, WT, or Y148D variant XPA after treatment with 100 nM bortezomib for indicated times (n = 3). β-tubulin included as loading control. Band intensities were calculated using FIJI. Normalized XPA intensity determined by first normalizing the values for each XPA lane to the corresponding β-tubulin signal and then back to that for the corresponding XPA protein at each 0 hour timepoint. Mean normalized XPA band intensities with standard deviation for 3 independent experiments shown in parentheses. ***E***, Representative Western blot images from analysis of stable XP2OS cells expressing mock XPA-null, WT, or Y148D variant XPA after treatment with 100 nM bortezomib for indicated times (n = 3). Poly-ubiquitin K48 included as UPS inhibition control. β-tubulin included as loading control. Band intensities were calculated using FIJI. Normalized XPA intensity determined by first normalizing the values for each XPA lane to the corresponding β-tubulin signal and then back to that for the corresponding XPA protein at each 0 hour timepoint.

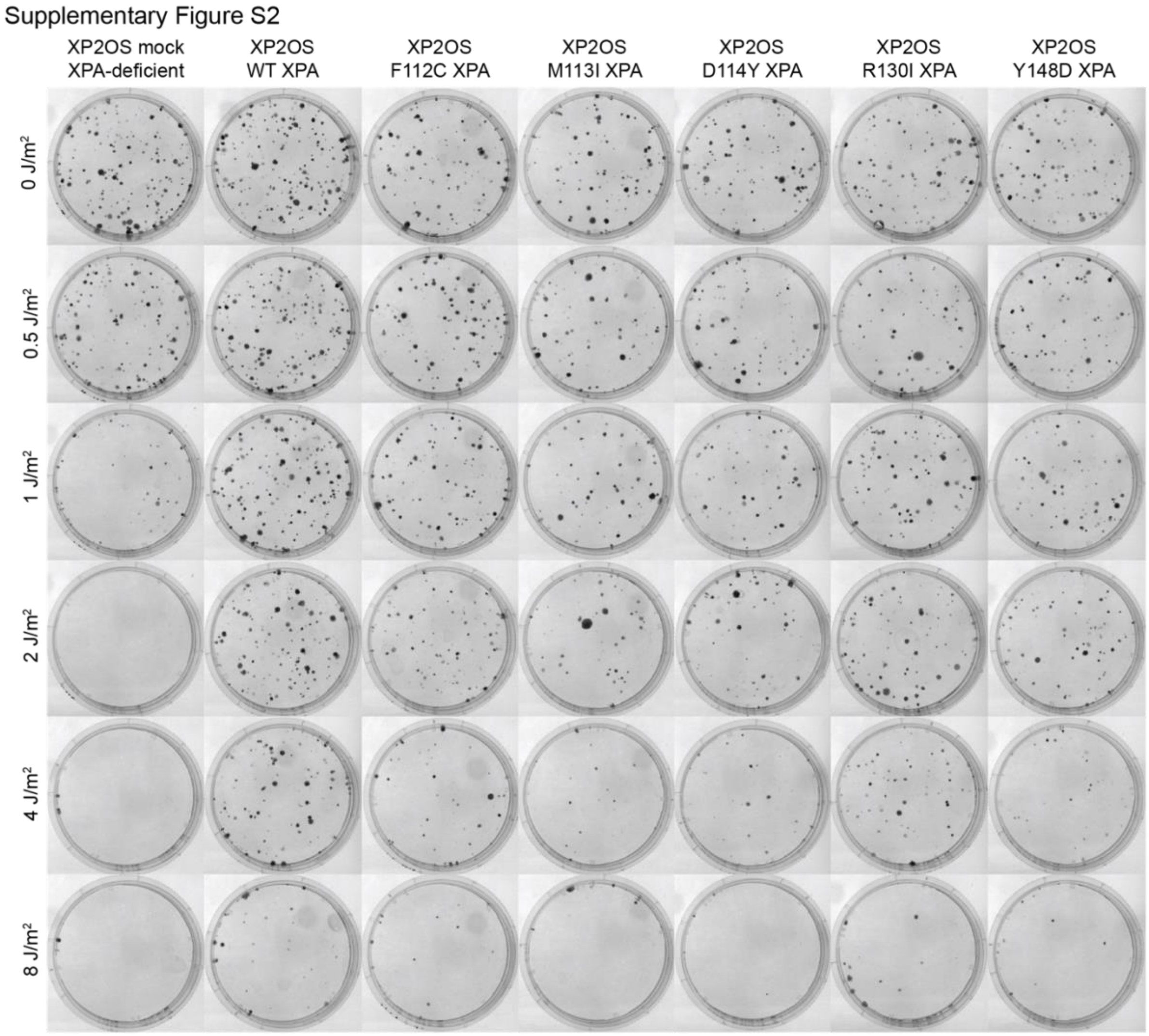


**Supplementary Figure S2. Representative images of UV clonogenic survival assay in stable XP2OS cells.** Images of crystal violet-stained cell colonies in 6 cm dishes, formed by stable XP2OS cell lines after exposure to UV at indicated doses and growth for 14 days.

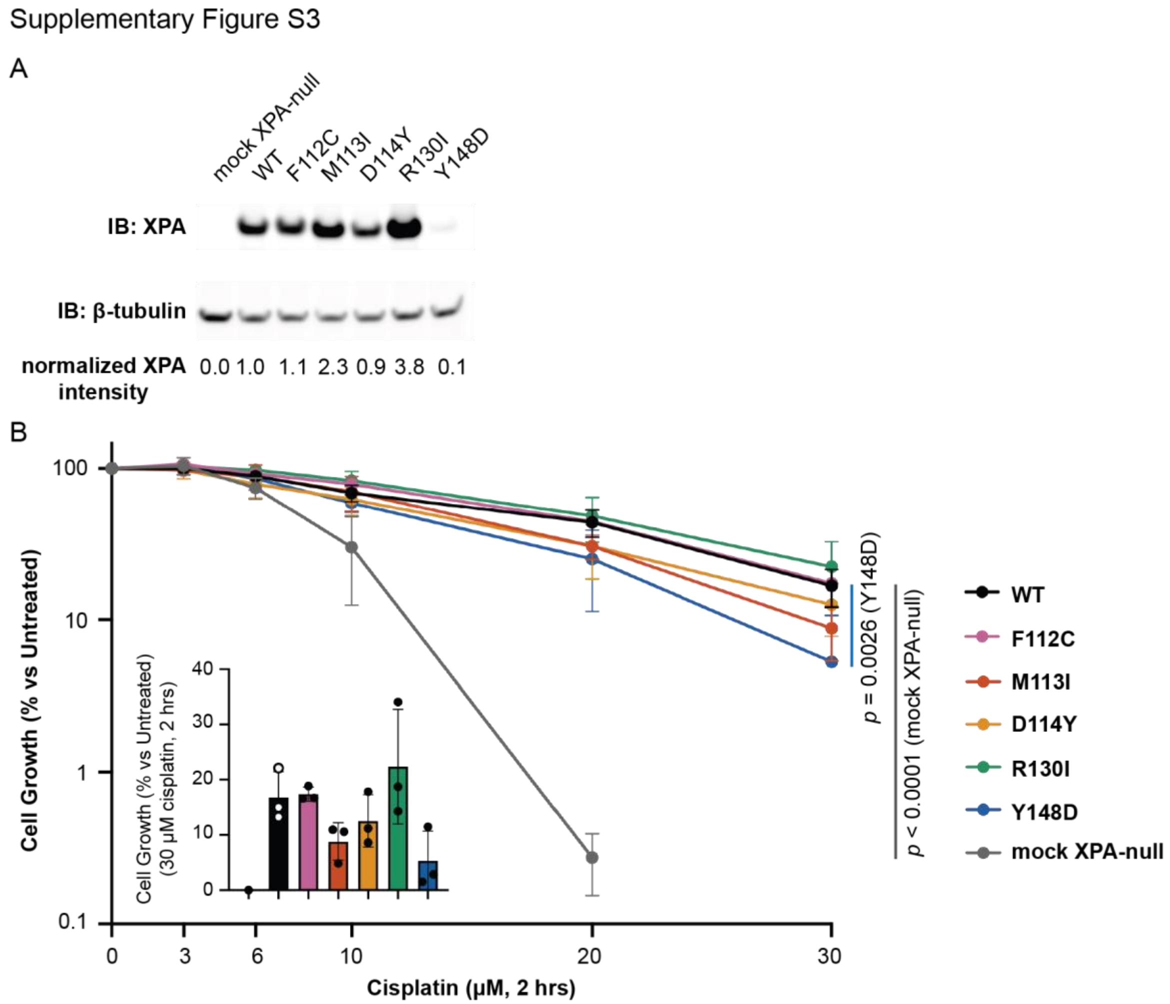


**Supplementary Figure S3. A second set of cells stably expressing NER-deficient XPA tumor variants has increased sensitivity to cisplatin.** ***A***, Western blot analysis of stable mock XPA-null, wild-type (WT) or variant XPA overexpression levels in XP patient-derived dermal fibroblast XP2OS cells transduced with lentivirus. β-tubulin included as loading control. Band intensities were calculated using FIJI. Normalized XPA intensity determined by first normalizing the values for each XPA lane to the corresponding β-tubulin signal and then back to that for WT XPA. ***B***, SRB assay in stable XP2OS cell lines after exposure to cisplatin for two hours at indicated doses and growth for five days (n = 3). Cells were stained with 0.4% SRB and absorbance at 490 nm measured. Percent untreated cell growth was determined for each cell line relative to untreated. $IC_{50}$ values [95% CI] determined using lines of best fit from nonlinear regression analysis in GraphPad Prism ([inhibitor] vs. response – variable slope (four parameters equation): 9 μM [7 – 10 μM] (mock XPA-null), 18 μM [15 – 20 μM] (WT), 19 μM [16 – 22 μM] (F112C), 15 μM [13 – 18 μM] (M113I), 14 μM [12 – 16 μM] (D114Y), 21 μM [17 – 24 μM] (R130I) and 14 μM [11 – 15 μM] (Y148D). EC50 shift, X is concentration equation used to compare the $IC_{50}$ of each line of best fit to that of the WT control, indicating comparisons to WT with $p < 0.05$. Inset showing mean cell growth for each cell line at 30 μM cisplatin.

Supplementary Figure S4

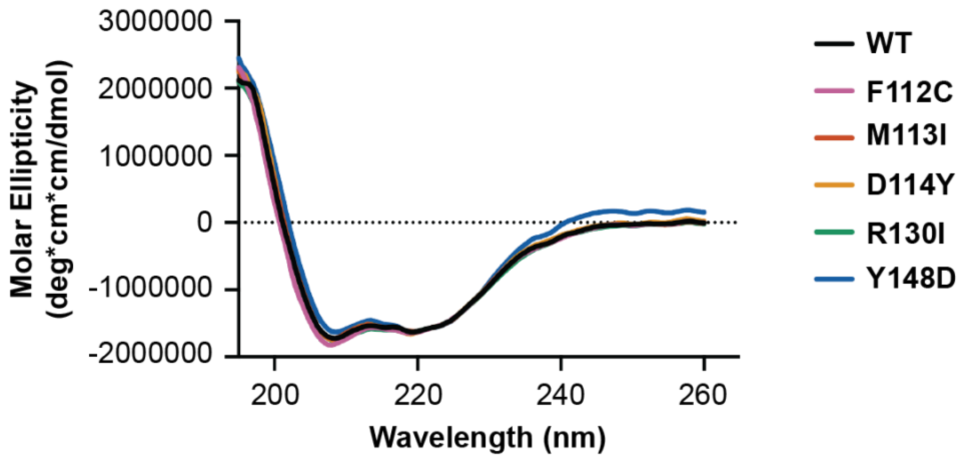


**Supplementary Figure S4. Global secondary structure of purified XPA variant proteins.** Circular dichroism spectra of purified recombinant WT or variant XPA DBD protein measured at 20 °C (n = 3). Each curve represents the average of three smoothed measurements. Ellipticities for the buffer and cuvette alone were subtracted from each measurement. Data were scaled to the WT spectrum at 222 nm.

Supplementary Figure S5

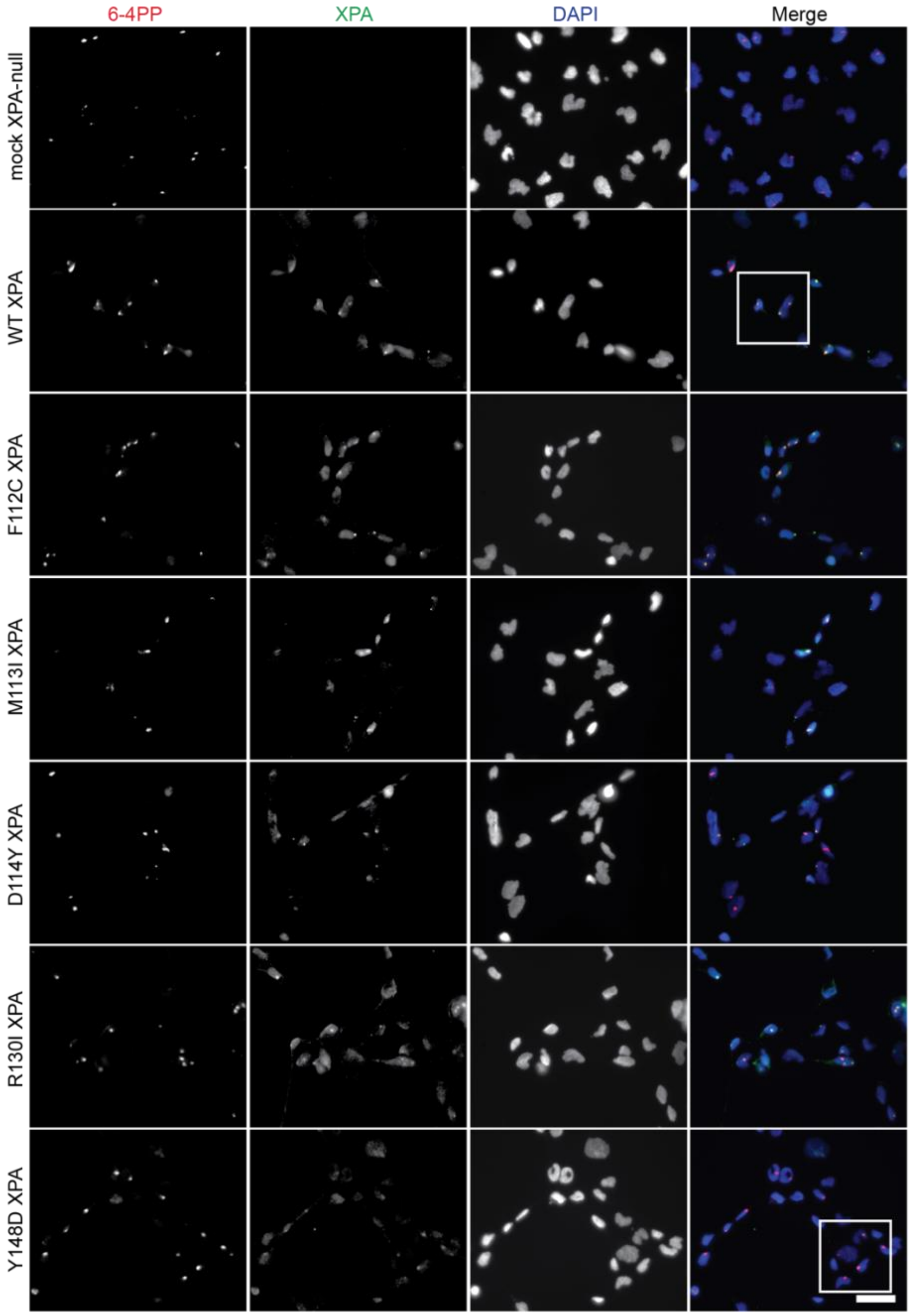


**Supplementary Figure S5. Representative images of XPA variant co-localization with UV damage in stable XP2OS cells.** Local UV irradiation and immunofluorescence assay in stable XP2OS cell lines. Cells were exposed to 120 J/$m^2$ UV irradiation through polycarbonate isopore membranes with 5 $\mu$m pores and XPA and 6-4PP damage foci were detected using immunofluorescence after 30 minutes incubation. Scale bar 40 $\mu$m. White boxes indicate crop for enlarged images shown in **Figure 6**.

**Supplementary Table 1. Mutagenesis primers to generate XPA variants in expression vectors.**

| Variant | Forward primer (5'-3') | Reverse primer (5'-3') |
|---|---|---|
| F112C | GGGAAAGAATgTATGGATTCTTATC | ACATTCTTCGCATATTACATAATC |
| M113I | AAGAATTTATtGATTCTTATCTTATGAACCAC | TCCCACATTCTTCGCATATTAC |
| D114Y | AGAATTTATGtATTCTTATCTTATGAACCAC | TTCCCACATTCTTCGCATATTAC |
| R130I | GATAACTGCAtcGATGCTGATGATAAAC | ACAAGTTGGCAAATCAAAG |
| Y148D | AAAACAAGAAgATCTTCTGAAAGAC | GCCTCTGTTTTGGTTATAAG |
